# Tuned normalization in perceptual decision-making circuits can explain seemingly suboptimal confidence behavior

**DOI:** 10.1101/558858

**Authors:** Brian Maniscalco, Brian Odegaard, Piercesare Grimaldi, Seong Hah Cho, Michele A. Basso, Hakwan Lau, Megan A. K. Peters

## Abstract

Current dominant views hold that perceptual confidence reflects the probability that a decision is correct. Although these views have enjoyed some empirical support, recent behavioral results indicate that confidence and the probability of being correct can be dissociated. An alternative hypothesis suggests that confidence instead reflects the magnitude of evidence in favor of a decision while being relatively insensitive to the evidence opposing the decision. We considered how this alternative hypothesis might be biologically instantiated by developing a simple leaky competing accumulator neural network model incorporating a known property of sensory neurons: tuned normalization. The key idea of the model is that each accumulator neuron’s normalization ‘tuning’ dictates its contribution to perceptual decisions versus confidence judgments. We demonstrate that this biologically plausible model can account for several counterintuitive findings reported in the literature, where confidence and decision accuracy were shown to dissociate -- and that the differential contribution a neuron makes to decisions versus confidence judgments based on its normalization tuning is vital to capturing some of these effects. One critical prediction of the model is that systematic variability in normalization tuning exists not only in sensory cortices but also in the decision-making circuitry. We tested and validated this prediction in macaque superior colliculus (SC; a region implicated in decision-making). The confirmation of this novel prediction provides direct support for our model. These findings suggest that the brain has developed and implements this alternative, heuristic theory of perceptual confidence computation by capitalizing on the diversity of neural resources available.

**Significance:** The dominant view of perceptual confidence proposes that confidence optimally reflects the probability that a decision is correct. But recent empirical evidence suggests that perceptual confidence exhibits a suboptimal ‘confirmation bias’, just as in human decision-making in general. We tested how this ‘bias’ might be neurally implemented by building a biologically plausible neural network model, and showed that the ‘bias’ emerges when each neuron’s degree of divisive normalization dictates how it drives decisions versus confidence judgments. We confirmed the model’s biological substrate using electrophysiological recordings in monkeys. These results challenge the dominant model, suggesting that the brain instead capitalizes on the diversity of available machinery (i.e., neuronal resources) to implement *heuristic* -- not optimal -- strategies to compute subjective confidence.

---

A dominant idea in the study of perceptual decision-making is that confidence judgments optimally reflect the probability that a decision is correct (Fetsch et al., 2014; Kiani et al., 2014; Pouget et al., 2016; Sanders et al., 2016; Zylberberg et al., 2016). Several models specifically stipulate that confidence is calculated via implementation of a diffusion framework: a decision is made when evidence for a decision reaches a certain threshold, and confidence reflects an optimal readout of the same information (Ratcliff and Rouder, 1998; Ratcliff and McKoon, 2008; Pleskac and Busemeyer, 2010; Tsetsos et al., 2012; Fetsch et al., 2014; Kiani et al., 2014; Zylberberg et al., 2016).

While this optimal ‘probability correct’ account of confidence has enjoyed significant empirical support, it seems difficult for it to account for cases where task performance and confidence dissociate (Rahnev et al., 2011, 2012b; Koizumi et al., 2015; Maniscalco et al., 2016; Samaha et al., 2016; Peters et al., 2017a; Odegaard et al., 2018). Seemingly suboptimal behaviors have also been observed in post-decisional perceptual judgments other than confidence (Stocker and Simoncelli, 2008; Luu and Stocker, 2018), leading these authors to hypothesize that these suboptimalities may stem from limitations on computational (i.e., neural) resources or a drive towards self-consistent behavior. One alternative theory of confidence, therefore, proposes that subjective confidence relies primarily on the magnitude of evidence supporting an observer’s decision, while ignoring or downplaying evidence supporting alternative, unchosen decisions (Zylberberg et al., 2012; Aitchison et al., 2015; Koizumi et al., 2015; Maniscalco et al., 2016; Samaha et al., 2016, 2017). In other words, to compute confidence the system uses a suboptimal heuristic that overly relies on decision-congruent evidence magnitude rather than optimal computations. Indeed, a recent study reported evidence for these decision-congruent evidence confidence computations using human intracranial electrocorticography (Peters et al., 2017b).

However, to date no biologically plausible mechanism has been proposed that might explain these dissociations between confidence and performance, or the decision-congruent confidence computations on which they seem to depend. We therefore developed a simple leaky competing accumulator network model (Usher and McClelland, 2001) to test a new hypothesis of how these computations might be implemented. This model extends previous work to incorporate a known property of perceptual circuitry: *tuned normalization* (Ni et al., 2012; Ruff et al., 2016; Verhoef and Maunsell, 2017), meaning each neuron is characterized by the specific degree to which it is normalized (i.e., inhibited) by surrounding network activity (Reynolds and Heeger, 2009; Carandini and Heeger, 2012), and specifically by units with opposing tuning preferences. We hypothesized that each neuron’s degree of tuned normalization dictates how it differentially participates in discrimination decisions versus confidence judgments. Specifically, we reasoned that highly normalized evidence accumulation neurons encode the balance of evidence for various perceptual interpretations (e.g. net evidence for leftwards or rightwards motion direction), and thus are ideally suited for making discrimination judgments. By contrast, less normalized evidence accumulation neurons encode evidence in favor of one perceptual interpretation (e.g. leftward motion) while ignoring evidence for alternative interpretations (e.g. rightward motion), and thus are ideally suited for implementing decision-congruent confidence computations. Therefore, the simple design principle that more normalized accumulator neurons drive decisions and less normalized accumulator neurons drive confidence may be sufficient to account for some of the most counterintuitive empirical findings on confidence in perceptual decision-making.

We tested key predictions of a *Differential Tuned Normalization* model instantiating this hypothesis using computational modeling and single neuron physiology. Model simulations show strong support for our hypothesis: the model reproduces multiple empirical findings when confidence is computed primarily from less normalized units, but not when computed primarily from more normalized units. Furthermore, we confirmed a critical prediction of the model -- that neurons in perceptual decision-making areas ought to exhibit tuned normalization -- using recordings from Rhesus macaque superior colliculus (a subcortical area involved in perceptual decision-making (Gold and Shadlen, 2000; Horwitz et al., 2004; Smith and Ratcliff, 2004; Kim and Basso, 2008). Our results suggest that tuned normalization may play a crucial role in how the brain differentially computes perceptual decisions and subjective confidence, revealing an important psychological function of this neuronal property.

## Materials and Methods

### 1. The *Differential Tuned Normalization* model

a leaky competing accumulator network with tuned normalization, where differently normalized units differentially contribute to perceptual decisions and confidence

To investigate how decision-congruent evidence might be biologically implemented, we began by considering known properties of perceptual decision-making circuitry. It is well known that divisive normalization is a canonical neural computation throughout the cortex (Simoncelli and Heeger, 1998; Reynolds and Heeger, 2009; Churchland, 2011; Ohshiro et al., 2011; Carandini and Heeger, 2012; Ling and Blake, 2012; Nassi et al., 2014). Further, it was recently reported that neurons in primary sensory areas exhibit *tuned normalization*, i.e. that each neuron possesses a unique, consistent degree of normalization: some neurons are very sensitive to activity of other units in the network (especially those which have different tuning preferences), while others operate more independently (Ni et al., 2012; Ruff et al., 2016; Verhoef and Maunsell, 2017). Previous implementations of evidence accumulation models for perceptual decisions have typically considered how a single level of normalization can account for behavioral data (Ratcliff and Rouder, 1998; Usher and McClelland, 2001; Ratcliff and McKoon, 2008; Tsetsos et al., 2012; Zylberberg et al., 2012, 2016; Fetsch et al., 2014; Kiani et al., 2014). However, we now know that a range of normalization tuning exists, at least in sensory cortices. We hypothesized that these neuron-by-neuron variations in normalization may reflect not noise or measurement error, but meaningful properties of the perceptual decision-making circuitry (Ni et al., 2012; Ruff et al., 2016; Verhoef and Maunsell, 2017). We refer to this model as the *Differential Tuned Normalization* model, or the *Tuned Normalization* model for short.

But *how* might tuned normalization be utilized in a behaving organism? To answer this, we should consider the tasks an organism must successfully execute in an ecologically valid environment. The ability to discriminate among multiple possible stimulus identities is certainly important, and for this type of task an observer ought to rely on a system that is able to average out noise, i.e. is less susceptible to random fluctuations in signal. Thus, for these discrimination-type tasks, a strong degree of lateral inhibition would be desirable, as it has been shown that neurons with stronger tuned normalization do exhibit weaker pairwise correlations (Ruff et al., 2016; Verhoef and Maunsell, 2017). But it is equally important that an organism also be able to detect a stimulus in the first place, regardless of its identity. For these detection-type tasks, such strong inhibition would be actually undesirable, as minute evidence amounts may be informative; therefore, weakly divisively normalized neurons ought to play a stronger role in detection-type tasks. As both of these task types are critical for an organism’s survival, it seems unlikely that a system would only be optimized for one or the other, which could in theory explain the presence of tuned normalization.

In light of this discussion, and of the empirically observed tuned normalization in cortical areas, a biologically plausible model of sensory evidence accumulation ought to implement more than one level of lateral inhibition and consider how such tuned normalization may affect a neuron’s role in the circuitry. Further, such stratification of tuned normalization could provide a neural mechanism to explain findings that confidence judgments rely on the magnitude of decision-congruent evidence (Zylberberg et al., 2012; Aitchison et al., 2015; Koizumi et al., 2015; Maniscalco et al., 2016; Peters et al., 2017b; Samaha et al., 2017). Specifically, the output of less normalized ‘detection’ neurons could be used to index decision-congruent evidence and therefore be used for confidence rating. This suggests that normalization tuning provides a biologically plausible mechanism to keep track of decision-congruent evidence independently of evidence favoring other possible choices by relying on the less normalized portion of the circuitry, while allowing the system to still capitalize on the beneficial consequences of divisive normalization by relying on the more normalized portion when discriminating among possible stimulus identities. We therefore hypothesized that normalization tuning might specifically dictate a neuron’s contribution to discrimination versus confidence judgments in decision-making circuitry.

We examined this hypothesis by incorporating tuned normalization (Ni et al., 2012; Ruff et al., 2016) into a leaky competing accumulator network (Usher and McClelland, 2001) (Figure 1). Intuitively, this network’s architecture can be summarized as follows. Accumulator units (with self-excitation and leak) tuned to varying stimulus alternatives accumulate momentary stimulus evidence and inhibit other units that have opposing tuning preferences. Units differ in the degree to which they are inhibited: the more normalized units are the ones that receive stronger inhibition. A discrimination decision is made when a linear combination of accumulator unit activity for a given tuning preference reaches a threshold level of evidence, and confidence is evaluated by reference to a linear combination of accumulator unit activity for the chosen stimulus alternative.

**Figure 1.**
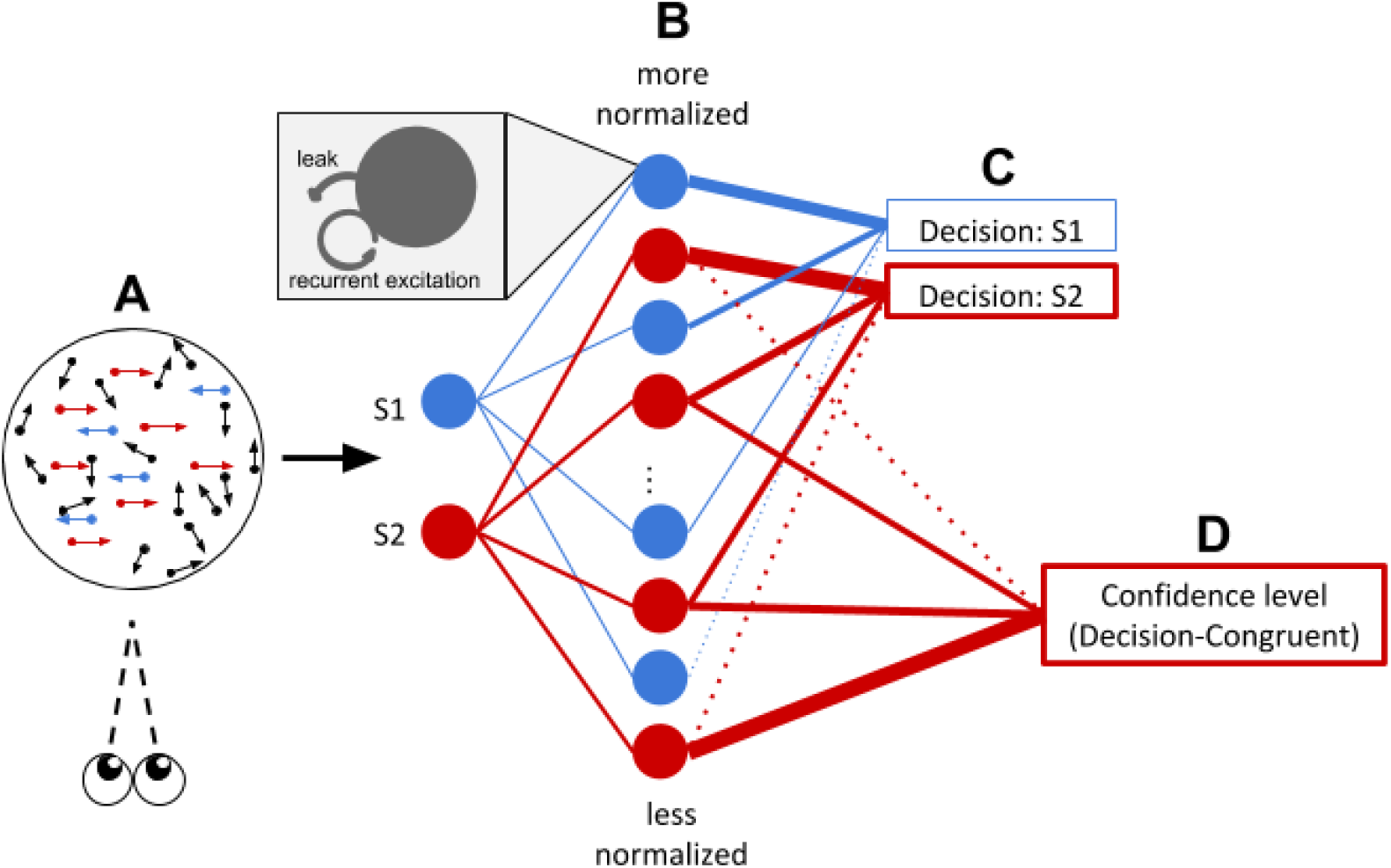
Tuned Normalization model architecture. (a) An observer views a stimulus that contains noisy information, with more information favoring S2 (red) than S1 (blue). (b) The network consists of accumulator units, each one of which has a specific and consistent degree of tuned normalization: more normalized units are more strongly inhibited by units that have opposing tuning preferences, whereas less normalized units operate relatively independently, neither inhibiting nor exciting other units in the circuit. Following convention (Usher and McClelland, 2001), all units also possess biologically plausible self-excitation and leak. (c) The *more* normalized units (upper) contribute more to the Decision variables favoring one or the other stimulus alternative, as these units’ activity represents the difference in accumulated evidence for the stimulus alternatives. A choice is made when one of these Decision variables reaches a boundary, indicating that the balance of evidence is sufficient for the observer to make a decision. In this example, the observer decides S2 (red), because the evidence for S2 was stronger than for S1. (d) Confidence is then read out as the magnitude of a Confidence variable for the winning alternative, relying more on activity of the *less* normalized units (lower), as these units’ activity represents the absolute magnitude of accumulated evidence for each stimulus alternative regardless of other information present in the stimulus. The model’s dynamics are fully mathematically described in Methods (Section S1).

Crucially, the weighting of accumulator units in these linear combinations depends on their level of normalization tuning, and this weighting differs for discrimination decisions and confidence ratings. More normalized units are weighted more heavily for discrimination decisions, since they effectively encode the balance of evidence for one stimulus alternative versus the others by virtue of their normalization. By contrast, less normalized units are weighted more heavily for confidence ratings, since they effectively encode a more faithful representation of the raw magnitude of evidence supporting each decision alternative, regardless of evidence supporting other possible decisions (Figure 1c & 1d). Details of model implementation and all simulations follows in the next sections.

#### 1.1 Accumulation of evidence

Extending previous work (Usher and McClelland, 2001), at each timestep we simulate the change in firing rate *dx* for each accumulation unit x with stimulus tuning preference *i* and normalization tuning level *k* (where *i* and *k* range from minimum values of 1 to maximum values of I and K, respectively) as

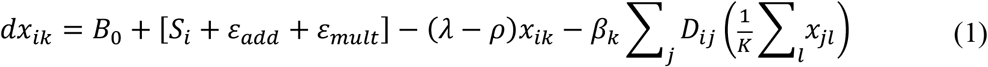

The components contributing to each unit’s momentary change in firing rate *dx* can be grouped as follows.

*Spontaneous firing rate*: *B_0_*

*B_0_* is a Poisson random variable such that *B_0_ ~ Poiss(b)*.

*Stimulus drive and associated noise: S_i_, ε_add_*, and *ε_mult_*

*S_i_* is the momentary magnitude of the stimulus drive to the unit with tuning preference *i*. The momentary stimulus drive is corrupted by two sources of noise: an additive Gaussian noise term *ε_add_* ~ *N*(0, *σ_add_*), and a multiplicative Gaussian noise term *ε_mult_ ~ N*(0, *σ_mult_*[*S_i_* + *ε_add_*]).

*Balance of self-excitation and leak*: – (*λ* – *ρ*)*X_ik_*

*ρ* is self-excitation and *λ* is leak. When *λ* > *ρ*, the accumulator unit “leaks” firing rate to a degree proportional to its current firing rate *x*.

*Lateral inhibition*: 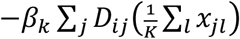

The term for lateral inhibition can be decomposed into two components.

1. Whole-population activation of inhibitory interneurons: 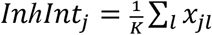

Every accumulator unit with tuning preference *j* and normalization tuning level *l* activates an inhibitory interneuron with the same tuning preference *j, Inhlnt_j_*, to a degree proportional to its current firing rate *X_jl_*. (Subscripts *j* and *l* are used here rather than *i* and *k* because the units subscripted with *j* and *l* are summed over the whole population of neurons in determining the effect of the population on a single unit with particular tuning preference *i* and normalization tuning level *k*, i.e. in determining *dx_ik_*). The summed activation across all normalization tuning levels *l* is divided by the overall number of tuning levels implemented in the simulation *K*. Thus, *Inhlnt_j_* is effectively an average of the activity of all accumulator units with tuning preference *j* computed across all normalization tuning levels *l*.

2. Inhibitory interneuron inhibition of single units: *Inh_X_ik_* = –*β_k_ Σ_j_D_ij_Inhlnt_j_*

Each inhibitory interneuron in turn inhibits each accumulator unit. The degree of inhibition depends on the dissimilarity in their respective tuning preferences *j* and *i*, according to the equation

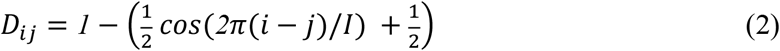

where I is the number of tuning preferences implemented in the simulation. *D_ij_* is thus a sinusoidal function varying between 0 and 1 that represents the dissimilarity in tuning preference between the accumulator unit with tuning preference *i* and the inhibitory interneuron with tuning preference *j* (Deneve et al., 1999). Tuning preference dissimilarity is maximal when | (*i* – *j*) / I | = ½ (i.e. when the accumulator unit and inhibitory interneuron have diametrically opposing tuning preferences, such as “motion left” vs “motion right”) and minimal when *i* = *j* (i.e. when they have the same tuning preference).

Inhibitory interneuron inhibition of accumulator units is further modulated by *β_k_*. The magnitude of *β_k_* is inversely proportional to normalization tuning level *k*, from a maximum value of 1 at *k* = 1 (full normalization tuning) to a minimum value of 0 at *k* = *K* (no normalization):

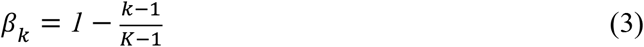

When *β_k_* = 0, a unit receives no lateral inhibition and is thus not normalized. When *β_k_* = 1, a unit is “fully normalized” in the sense that it receives momentary inhibition from the average activity of all units with opposite tuning preference (for which *D_ij_* = 1). Intermediate levels of *β_k_* reflect intermediate levels of normalization.

A schematic of the model’s implementation of lateral inhibition is presented in Figure 2.

**Figure 2.**
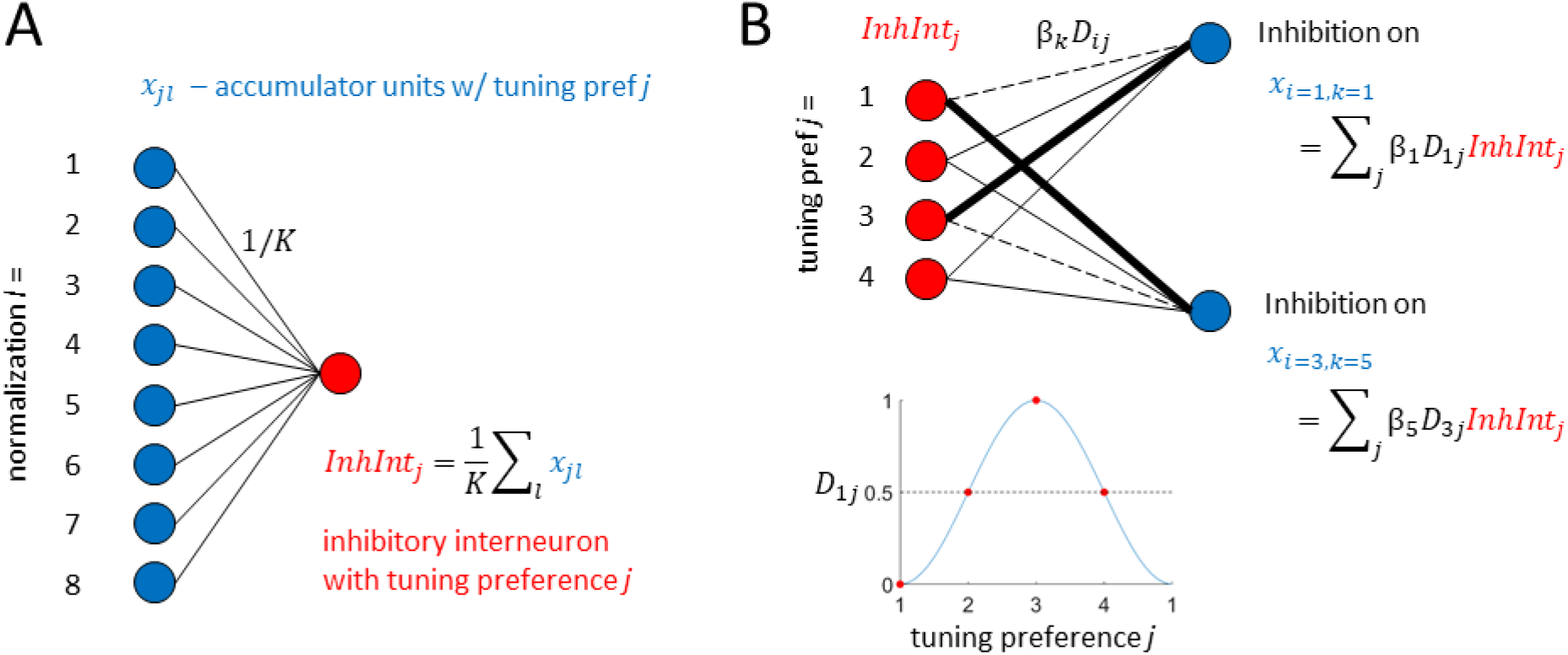
Schematic of lateral inhibition in the model. (A) *Inhibitory interneuron units.* There are accumulator units with tuning preference *j* at each level of normalization tuning *l*. Activity in these units is integrated into a single inhibitory interneuron unit with the same tuning preference, *InhIntj*, where the weighting factor on each unit is set to a constant 1/K, where K is the number of normalization tuning levels (8 in this example). (B) *Inhibition of accumulator units according to stimulus tuning preference dissimilarity.* Inhibitory interneuron pools *InhIntj* inhibit accumulator units *x_ik_* as a sinusoidal function of the similarity in tuning preferences *i* and *j*, *D_ij_*. For instance, for a set of I = 4 circularly arranged tuning preferences (corresponding e.g. to motion directions left, up, right, down), an accumulator unit with tuning preference *i* = 1 receives weakest inhibition from inhibitory interneuron pools with the same tuning preference *j* = 1, and strongest inhibition from inhibitory interneuron pools with the opposite tuning preference *j* = 3. Here we illustrate an example using I = 4, although in the actual simulations we set I = 2 for simplicity, corresponding to diametrically opposed stimulus properties, e.g. motion directions left vs. right. The overall inhibition strength *β_k_* depends on normalization tuning level *k*, with more normalized units receiving stronger inhibition.

#### 1.2. Decisions and Confidence judgments

We can now examine how the network described above might perform a perceptual decision-making task and then provide a confidence judgment about its decision. Assuming a classic two-choice discrimination task (e.g., “Is the dot motion predominantly leftward or rightward?”), we can simulate units that have various tuning preferences and varying levels of normalization tuning. For the network to perform this two-choice discrimination task, we will assign the first stimulus alternative (e.g., “leftward motion”) as *S*_1_, and the second stimulus alternative (e.g., “rightward motion”) as *S*_2_.

The accumulating variables for the Decision between these two alternatives and the Confidence judgment about this decision are then defined as functions of weighted linear sums of network activity. The accumulated evidence E in favor of selecting stimulus alternative Si at time t is defined as

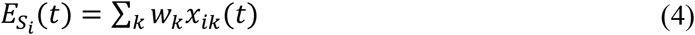

where *x_ik_*(*t*) refers to the activity at time *t* of the unit with tuning preference *i* (i.e., centered on stimulus *S_i_*) and normalization tuning level *k*. The weight for units with normalization level *k*, *w_k_*, is defined as being exponentially proportional to those units’ level of normalization tuning, i.e.

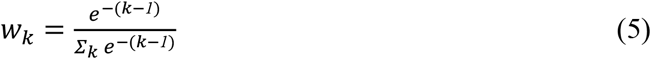

where the *w_k_* are normalized so that all weights sum to 1.

The two-choice discrimination decision is then made when the value of the decision variable E for one of the stimuli alternatives reaches a threshold *T*, i.e. when for either *S_i_*

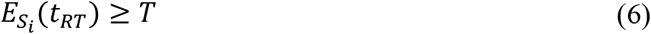

with *t_RT_* referring to the first time *t* at which Equation 6 is satisfied. The chosen stimulus is defined as the stimulus that produced the ‘winning’ decision variable value at this ‘reaction time’ *t_RT_*, i.e., *S_selected_* = *argmax_s_*(*E_s_*(*t_RT_*).

At the time when the perceptual decision is reached, confidence is read out as the accumulated confidence variable *C favoring the selected stimulus alternative*, i.e. the Decision-Congruent evidence, where the value of C at time *t* for stimulus alternative *S_selected_* is defined similarly to Equation 4 as

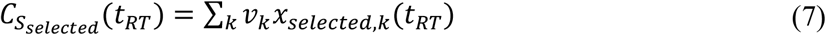

where the subscript changes from *x_ik_*(t) in Equation S4 to *x_sekcted,k_*(t_RT_) to reflect that confidence depends only on units whose tuning preferences are congruent with the selected stimulus, and the specification of t_RT_ indicates that confidence is reported based on the state of accumulator units at the time when the stimulus is chosen.

Crucially, whereas stimulus evidence E is computed using weights *w_k_* ∝ × *e^−k^*, confidence C is computed using alternative weights *v_k_* where

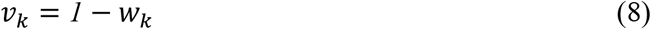

Thus, while the contributions of accumulator units to stimulus evidence E are directly proportional to their degree of normalization tuning (more normalization → more weight), the contributions of accumulator units to confidence C are inversely proportional to their degree of normalization tuning (more normalization → less weight).

To assess the importance of using the *v_k_* weights in Equation 8 for capturing empirical dissociations in confidence, in every trial of every simulation we also computed a secondary confidence variable C* using the same *w_k_* weights used to compute stimulus evidence E:

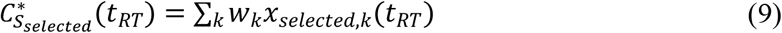

We used the confidence variable C* to compute confidence for all Null model simulations reported in the manuscript. The fact that the model was able to capture empirical patterns when computing confidence from C but not C* indicates that usage of the *v_k_* = 1 – *w_k_* weights in C was crucial for capturing those empirical patterns.

Because the stimulus evidence variable E relies heavily on more normalized units, it can be thought of as a readout of the balance of evidence for *S*_1_ versus *S*_2_. In contrast, because of its heavier reliance on less normalized units, the confidence variable C provides a better measure of exactly how much information has been collected for each stimulus alternative individually, regardless of the accumulated information for the opposing stimulus. In this way, the confidence variable C provides more information about the absolute magnitude of information in the environment than the stimulus evidence variable E and the alternative confidence variable C*.

Finally, to convert the continuous C and C* variables to discrete confidence ratings on an ordinal rating scale with N_ratings_ possible options, we defined a threshold parameter *U_r_* where the subscript *r* ranges from 1 to N_ratings_ – 1. The conversion of C to a discrete rating R can then be computed as

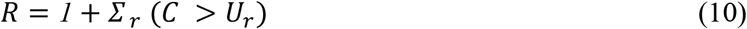

where (*C* > *U_r_*) is a logical comparison evaluating to 1 if the inequality is true and 0 otherwise. Thus, R is a simple count of how many of the confidence thresholds *U_r_* are surpassed by C, with the constant 1 added to set 1 as the minimum confidence rating value.

#### 1.3. Parameter setting and specifics of model implementation

Rather than attempt to fit the many parameters of the model to each individual data set, we adopted a simplified approach in which we fixed most model parameters for all simulations. Model parameters used for different simulations therefore differed only by (1) parameter values used for stimulus strength, (2) a single parameter coding across-condition differences in evidence accumulation noise ((Rahnev et al., 2012b) simulation; Figure 3) or stimulus volatility ((Zylberberg et al., 2016) simulation; Figure 4), and (3) confidence thresholds *U_r_*. Conceptually, the simulations were therefore set up as if the same observer with the same neural network were performing separate experiments with separate stimulus inputs and experimental manipulations. This approach was in line with our principle aim for this modeling project -- to capture qualitative patterns in behavioral effects reported in the literature and thereby provide a proof of concept that those patterns can be explained by the biologically plausible mechanisms of the Tuned Normalization model. This simplifying approach also helps prevent overfitting and eases comparison of modeling results across simulations. A full list of parameter values used in all simulations is listed in Table 1.

**Figure 3.**
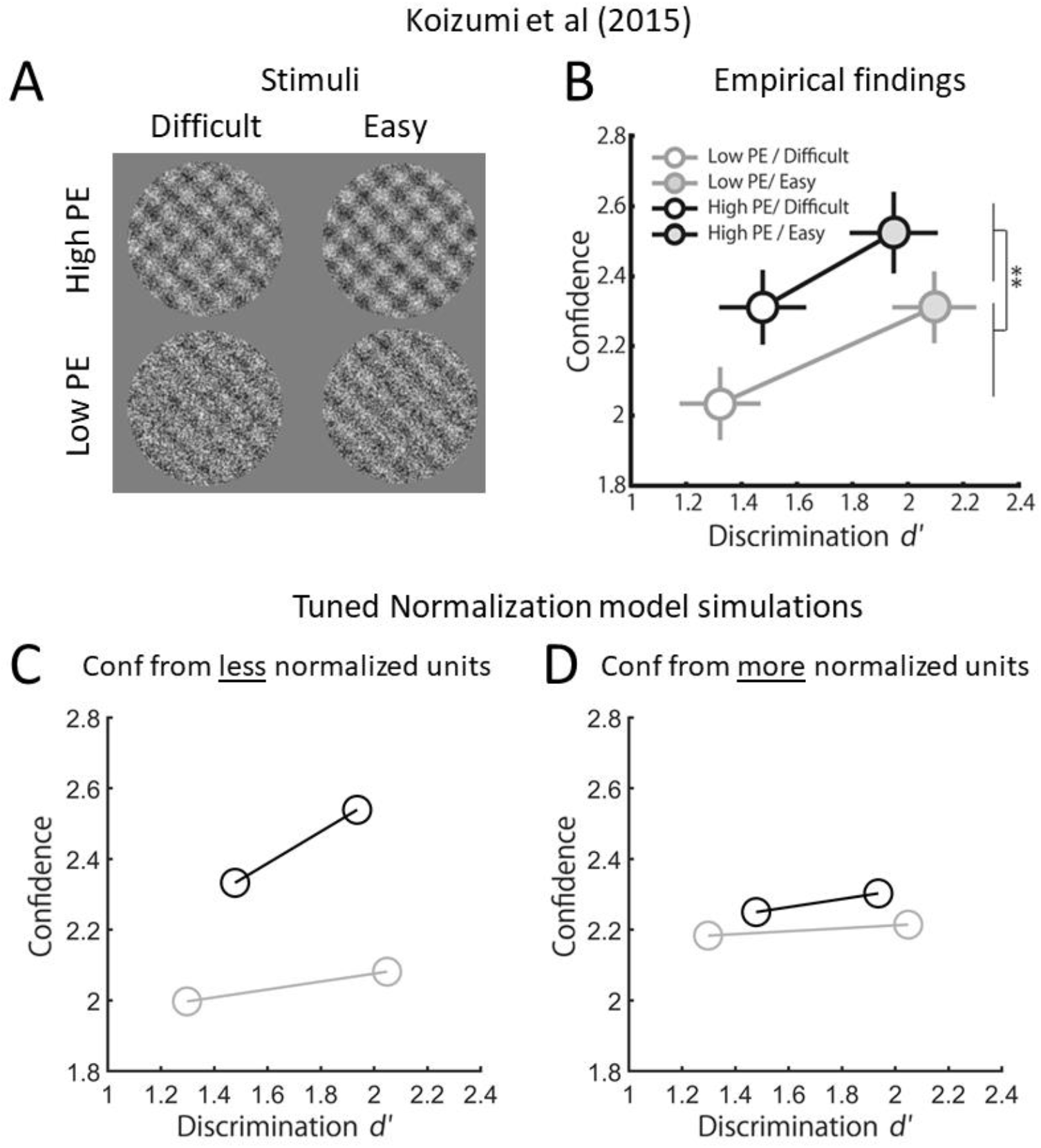
The Tuned Normalization model can account for effects of Positive and Negative Evidence on confidence. (a) (Koizumi et al., 2015) presented subjects with superimposed left-tilting and right-tilting gratings, where the task was to determine the predominant (highest contrast) tilt in the stimulus. Positive Evidence (PE) corresponded to the contrast of the target (higher-contrast) grating and Negative Evidence (NE) corresponded to the contrast of the distractor (lower-contrast) grating. Two levels of task difficulty were created by adjusting the PE / NE ratio. High and low PE conditions were created for each difficulty level. (b) Koizumi et al’s results showed that confidence in High Positive Evidence (High PE) conditions was higher than in Low Evidence (Low PE) conditions, and higher for Easy than for Difficult trials. (b) The *Tuned Normalization* model reproduces the observation that High Positive Evidence leads to higher confidence than Low Positive Evidence, and that Easy conditions produce higher confidence than Difficult conditions. (c) Confidence effects are almost completely abolished in control simulations in which more normalized units are weighted more heavily for computing confidence, suggesting that the result in (b) critically depends on confidence being computed primarily from less normalized units. Cohen’s d for confidence effects in the main model simulations were about 5 times greater than in the control simulations, and more closely matched the empirical effect sizes reported by Koizumi et al. (2015).

**Figure 4.**
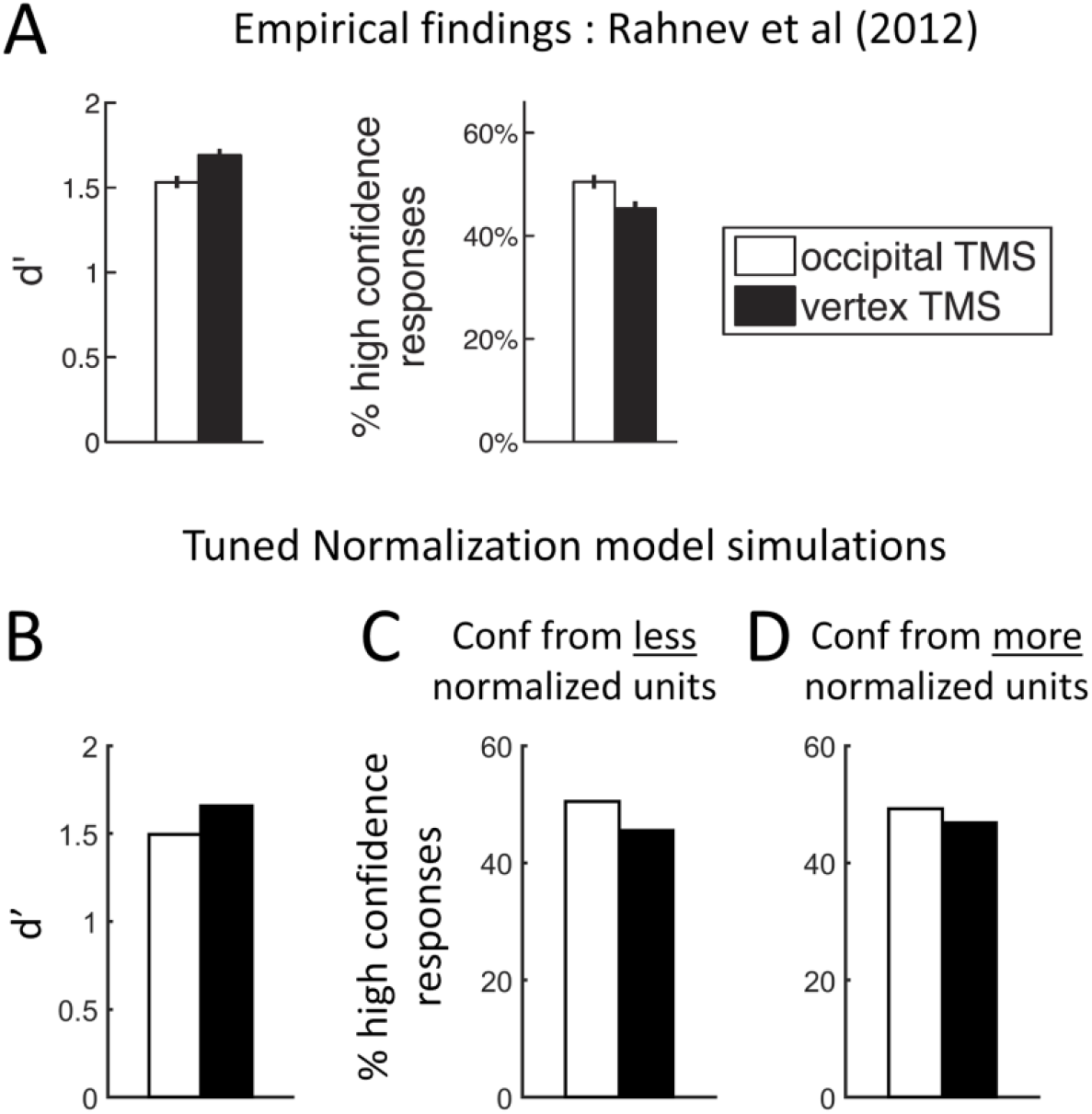
The Tuned Normalization model can account for effects of evidence accumulation noise on task performance and confidence. (a) Rahnev and colleagues (Rahnev et al., 2012) showed that occipital TMS decreased performance but increased confidence in a visual discrimination task, in line with other studies showing similar effects as a result of TMS (Rahnev et al., 2012b; Peters et al., 2017a), inattention (Rahnev et al., 2011), and microstimulation (Fetsch et al., 2014). (b, c) The Tuned Normalization model reproduces the observation that added evidence accumulation noise decreases task performance but increases confidence. Here we show a model fit to Rahnev et al.’s representative findings, thereby establishing the model’s ability to account for the general pattern in all the previously mentioned studies in which increasing noise decreases task performance while increasing confidence. (d) Confidence effects are almost completely abolished in control simulations in which more normalized units are weighted more heavily for computing confidence, suggesting that the result in (b) critically depends on confidence being computed primarily from less normalized units. Cohen’s d for confidence effects in the main model simulations were twice as large as in the control simulations across several levels of simulated noise, including larger noise levels that yielded larger behavioral effects than in the Rahnev et al. data set (see Results; extra simulations not shown).

**Table 1.**
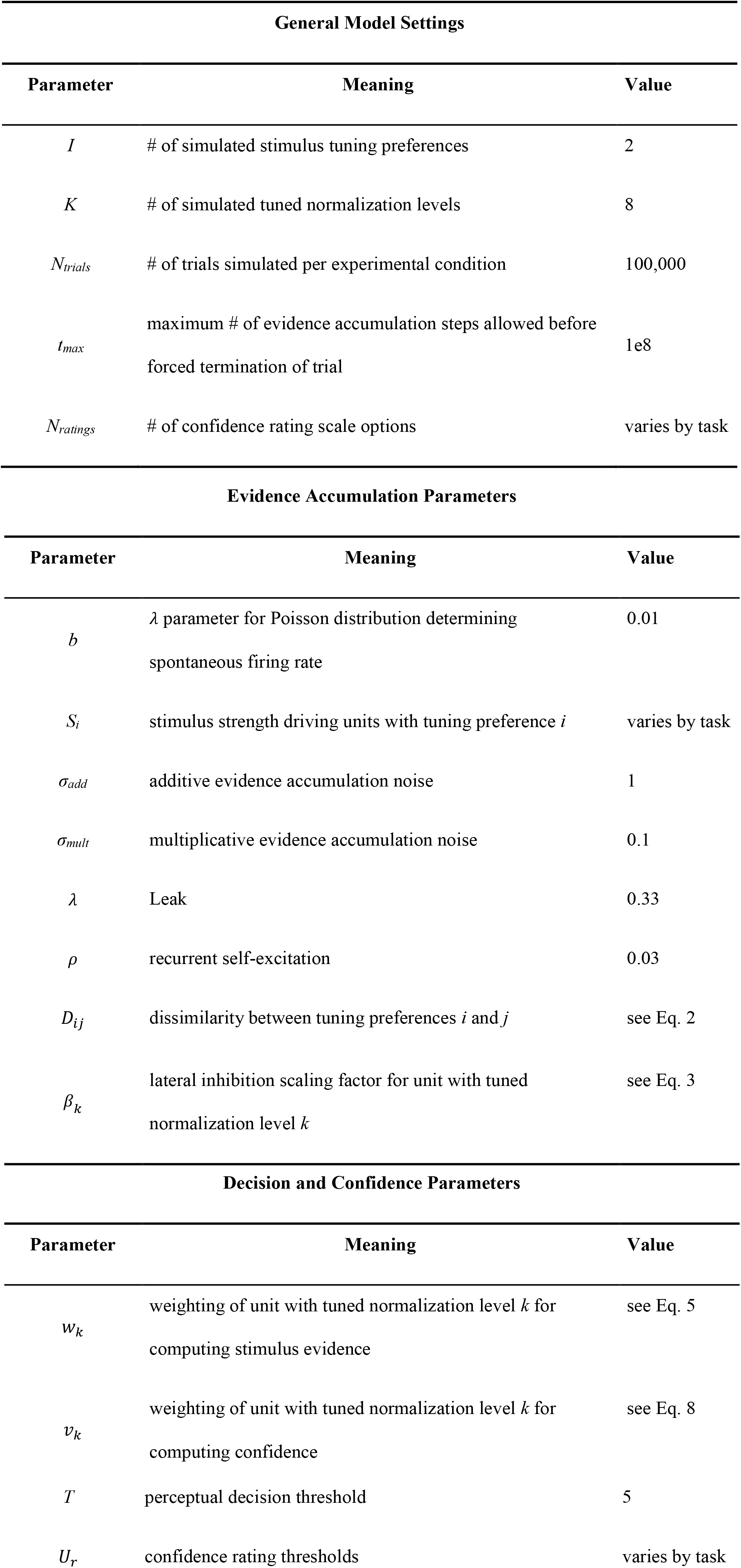
Model parameters fixed across all simulations.

| General Model Settings |  |  |
| --- | --- | --- |
| Parameter | Meaning | Value |
| $I$ | # of simulated stimulus tuning preferences | 2 |
| $K$ | # of simulated tuned normalization levels | 8 |
| $N_{\text{trials}}$ | # of trials simulated per experimental condition | 100,000 |
| $t_{\text{max}}$ | maximum # of evidence accumulation steps allowed before forced termination of trial | 1e8 |
| $N_{\text{ratings}}$ | # of confidence rating scale options | varies by task |

Evidence Accumulation Parameters
| Parameter | Meaning | Value |
| --- | --- | --- |
| $b$ | $\lambda$ parameter for Poisson distribution determining spontaneous firing rate | 0.01 |
| $S_i$ | stimulus strength driving units with tuning preference $i$ | varies by task |
| $\sigma_{add}$ | additive evidence accumulation noise | 1 |
| $\sigma_{mult}$ | multiplicative evidence accumulation noise | 0.1 |
| $\lambda$ | Leak | 0.33 |
| $\rho$ | recurrent self-excitation | 0.03 |
| $D_{ij}$ | dissimilarity between tuning preferences $i$ and $j$ | see Eq. 2 |
| $\beta_k$ | lateral inhibition scaling factor for unit with tuned normalization level $k$ | see Eq. 3 |

Table 1. Model parameters fixed across all simulations.
| Parameter | Meaning | Value |
| --- | --- | --- |
| $w_k$ | weighting of unit with tuned normalization level $k$ for computing stimulus evidence | see Eq. 5 |
| $v_k$ | weighting of unit with tuned normalization level $k$ for computing confidence | see Eq. 8 |
| $T$ | perceptual decision threshold | 5 |
| $U_r$ | confidence rating thresholds | varies by task |

##### General Model Settings

We chose K = 8 to model a range of normalization tuning levels, and I = 2 for simplicity. We verified that model results were reproducible using I = 4, K = 4, and K = 16. We set N_trials_ and t_max_ to high values to ensure the robustness of simulation output and to mitigate the effect of premature termination of evidence accumulation on model output. N_ratings_ was varied to match the confidence rating scale used in the empirical data being simulated.

##### Evidence Accumulation and Decision Parameters

The leak and recurrent self-excitation parameters were set according to values established by (Usher and McClelland, 2001). All other parameter values were determined by the authors, as follows.

We set the values of *D_ij_* and *β_k_* on *a priori* grounds. Both variables range from 0 to 1 and so represent simple scaling variables. *D_ij_* scales the magnitude of inhibition exerted on an accumulator unit *x_ik_* by an inhibitory interneuron according to the dissimilarity of their tuning preferences *i* and *j*, and *β_k_* scales this inhibition further according to *x_ik_*‘s normalization tuning level *k*.

The values of *w_k_* and *v_k_* were similarly set on *a priori* grounds. *w_k_* was chosen to be exponentially proportional to *k* as a mathematically convenient way to capture the idea that evidence accumulation is dominated by highly normalized units, while still allowing for less normalized units to have some influence. *v_k_* was defined as 1 − *w_k_* as a mathematically simple way to capture the complementary idea that confidence rating is dominated by less normalized units, while still allowing for highly normalized units to have some influence.

The remaining parameters *b, σ_add_, σ_mult_*, and *T* had no clear empirical or *a priori* motivations for defining parameter values. (We discuss *U_r_*, which transforms continuous confidence values to discrete confidence ratings, below.) Therefore, we simply set *σ_add_* = 1 arbitrarily and manually searched for values of the remaining parameters that, given the fixed values for the parameters already mentioned, would reliably yield intermediate levels of task performance (e.g. *d*’ ≈ 2) within a reasonable number of accumulation steps (~1000) for ‘mock’ simulations not directly based on any of the empirical studies being simulated. The model produced reasonable output across a wide range of parameter settings, and thus does not depend critically on the specific parameter values chosen here.

With the main model parameters thus determined, values for the stimulus strength parameter *S_i_* in individual simulations were determined by manual parameter search yielding task performance (*d*’ or percent correct) closely resembling empirical data. Importantly, values for stimulus strength parameters in each simulation were not altered after matching task performance to the relevant empirical data; in other words, confidence effects were merely ‘read out’ after task performance was best matched to the intended behavioral effect.

Confidence thresholds *U_r_* were set after running the simulation and obtaining the distribution of values of the confidence variable C across all simulated trials. *U_r_* was initialized as an evenly spaced set of quantile values on the distribution of C across all trials, and then manually adjusted to yield mean confidence rating output resembling the empirical data to be fitted. Thus, the same confidence thresholds *U_r_* were applied to all trials of all conditions within a given simulation, and these thresholds were chosen so as to approximately evenly span the full distribution of continuous confidence values produced by the simulation. A similar procedure was performed separately on the distribution of C* across all trials to determine confidence ratings based upon C*.

Because the same *U_r_* were applied to all trials of a given simulation, any between-condition differences in confidence arose purely from the influence of network activity on C (or C*), independently of the choice of *U_r_*. Changing confidence thresholds mainly affected mean confidence values, while retaining similar standard deviations and between-condition differences. Therefore, the effect size of differences in confidence between conditions was robust against choice of confidence thresholds.

### 2. Task-specific model settings

#### 2.1. Controlling positive and negative evidence to yield equal performance and unequal confidence

In Experiments 1A and 2A of (Koizumi et al., 2015)), subjects performed a grating tilt discrimination task. However, stimuli were actually composed of two superimposed gratings tilting in opposite directions, one with higher contrast (“Positive Evidence” or “PE” for short) and one with lower contrast (“Negative Evidence” or “NE”) (Figure 3a). Subjects had to indicate whether the higher contrast grating was tilting left or right and rate confidence. The key experimental manipulation was the introduction of High PE and Low PE conditions, in which the contrast of the NE gratings was set to 0.7*(PE grating contrast) and 0.35*(PE grating contrast), respectively, and PE grating contrast was controlled by thresholding procedures to achieve a criterion level of task performance. The High PE and Low PE conditions yielded similar task performance, but confidence was higher for High PE stimuli. Koizumi et al. collected data for the High PE and Low PE conditions at two levels of task difficulty.

We modeled this task as follows. On every trial, the evidence accumulated for the stimulus alternatives *S*_1_ and *S*_2_ (corresponding here to left and right tilt) was driven by the corresponding PE and NE values, call them *S_PE_* and *S_NE_*. For half of all trials, we set *S*_1_ = *S_PE_* and *S*_2_ = *S_NE_*, and vice versa for the other half. As in Koizumi et al., we defined *S_NE_* = 0.7 * *S_PE_* in the High PE condition and *S_NE_* = 0.35 * *S_PE_* in the Low PE condition. PE values were chosen to yield simulated *d*’ values closely matching the empirical values (Table 2).

**Table 2.** Simulation-specific values used for simulating the results of (Koizumi et al., 2015). Values for *S_NE_* are presented as functions of *S_PE_* to emphasize that these were fixed rather than free parameters.

|  | Difficult |  | Easy |  |
| --- | --- | --- | --- | --- |
|  | Low PE | High PE | Low PE | High PE |
| $S_{PE}$ | 0.1382 | 0.3384 | 0.2231 | 0.4562 |
| $S_{NE}$ | $0.35 * S_{PE}$ | $0.7 * S_{PE}$ | $0.35 * S_{PE}$ | $0.7 * S_{PE}$ |

To convert the model’s continuous confidence variables C to confidence ratings, we set N_ratings_ = 4 to match the confidence scale used in Koizumi et al., and chose *U*_r_ values to yield simulated mean confidence values closely matching the empirical values. The *U*_r_ values thus chosen corresponded to the 0.3812, 0.5875, and 0.7937 quantiles of the distribution of C. For deriving confidence ratings from C*, we used the same quantile-based definition of *U*_r_ as applied to the distribution of C*.

For a similar demonstration of dissociating confidence from task performance by controlling positive evidence, see (Samaha et al., 2016).

#### 2.2. Introducing noise: TMS, inattention, or microstimulation

Our simulation capturing the effect of increased perceptual noise due to TMS, inattention, or microstimulation focused on the representative study of (Rahnev et al., 2012b), which is conceptually similar to (Fetsch et al., 2014), (Peters et al., 2017a), and (Rahnev et al., 2011) discrimination tasks. In (Rahnev et al., 2012b), subjects performed a visual discrimination task in a control condition (vertex TMS) and a condition introducing increased perceptual noise (occipital TMS). They found that with increased noise, task performance decreased but confidence increased.

We modeled this task as follows. On every trial, the evidence accumulated for the stimulus alternatives *S*_1_ and *S*_2_ (corresponding here to left and right tilt of a small bar) was driven by the corresponding stimulus-present and stimulus-absent values, call them *S_stim_* and *S_null_*. For half of all trials, we set *S*_1_ = *S_stim_* and *S*_2_ = *S_null_*, and vice versa for the other half. We set *S_null_* = 0 and chose the value for *S_stim_* that yielded a simulated *d*’ value closely matching the empirical value of *d*’ in the vertex (control) TMS condition of (Rahnev et al., 2012b). Then, holding *S_stim_* constant, we increased the value of the additive evidence accumulation noise parameter *σ_add_* so as to yield a simulated *d*’ value closely matching the empirical *d*’ in the occipital TMS condition. These parameter settings are summarized in Table 3.

**Table 3.** Simulation-specific parameter values used for simulating the results of (Rahnev et al., 2012b). The columns labeled Baseline and Added Noise correspond to the vertex TMS and occipital TMS conditions of Rahnev et al.’s study.

|  | Baseline | Added Noise |
| --- | --- | --- |
| $S_{null}$ | 0 | 0 |
| $S_{stim}$ | 0.115 | 0.115 |
| $\sigma_{add}$ | 1 | 1.05 |

To convert the model’s continuous confidence variable C to confidence ratings, we set N_ratings_ = 2 to match the confidence scale used in Rahnev et al., and chose *U*_r_ to yield simulated mean confidence in the vertex TMS condition closely matching the empirical value. The *U*_r_ value thus chosen corresponded to the 0.52 quantile of the distribution of C. For deriving confidence ratings from C*, we used the same quantile-based definition of *U*_r_ as applied to the distribution of C*.

#### 2.3. Stimulus volatility leads to equal performance but increasing confidence

In (Zylberberg et al., 2016), subjects performed a motion discrimination task at varying levels of motion coherence. In the Low Volatility condition, motion coherence was constant throughout a trial, whereas in the High Volatility condition, motion coherence changed over the course of a trial. Specifically, motion coherence on each frame was randomly drawn from a Gaussian distribution of possible values. Task performance was unaffected by stimulus volatility, but confidence at weak motion coherences was higher in the High Volatility condition.

We modeled this task as follows. On every trial, the evidence accumulated for the stimulus alternatives *S*_1_ and *S*_2_ (corresponding here to left and right motion direction) was driven by the corresponding stimulus-present and stimulus-absent values, call them *S_stim_* and *S_null_*. For half of all trials, we set *S*_1_ = *S_stim_* and *S*_2_ = *S_null_*, and vice versa for the other half. We set *S_null_* = 0 and chose the value for *S_stim_* that yielded simulated p(correct) values increasing from chance to strong performance (~90% correct), similar to empirical values in Zylberberg et al. (2016). Then, to simulate the High Volatility condition, we introduced a new model parameter, *σ*_stim_. At each time step of evidence accumulation, we defined the instantaneous value of stimulus strength as a random sample from a Gaussian distribution, 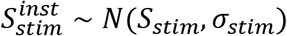. Following the methods of Zylberberg et al., in cases where 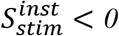, we set *S_stim_* = 0 and 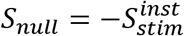 (corresponding to momentary stimulus evidence for the incorrect stimulus choice). Because the range of simulated confidence values was small compared to the empirical values in Zylberberg et al., we did not focus on achieving an exact fit to the data but simply selected a value for *σ_stim_* that could reproduce the qualitative patterns in the confidence data. These parameter settings are summarized in Table 4.

**Table 4.**
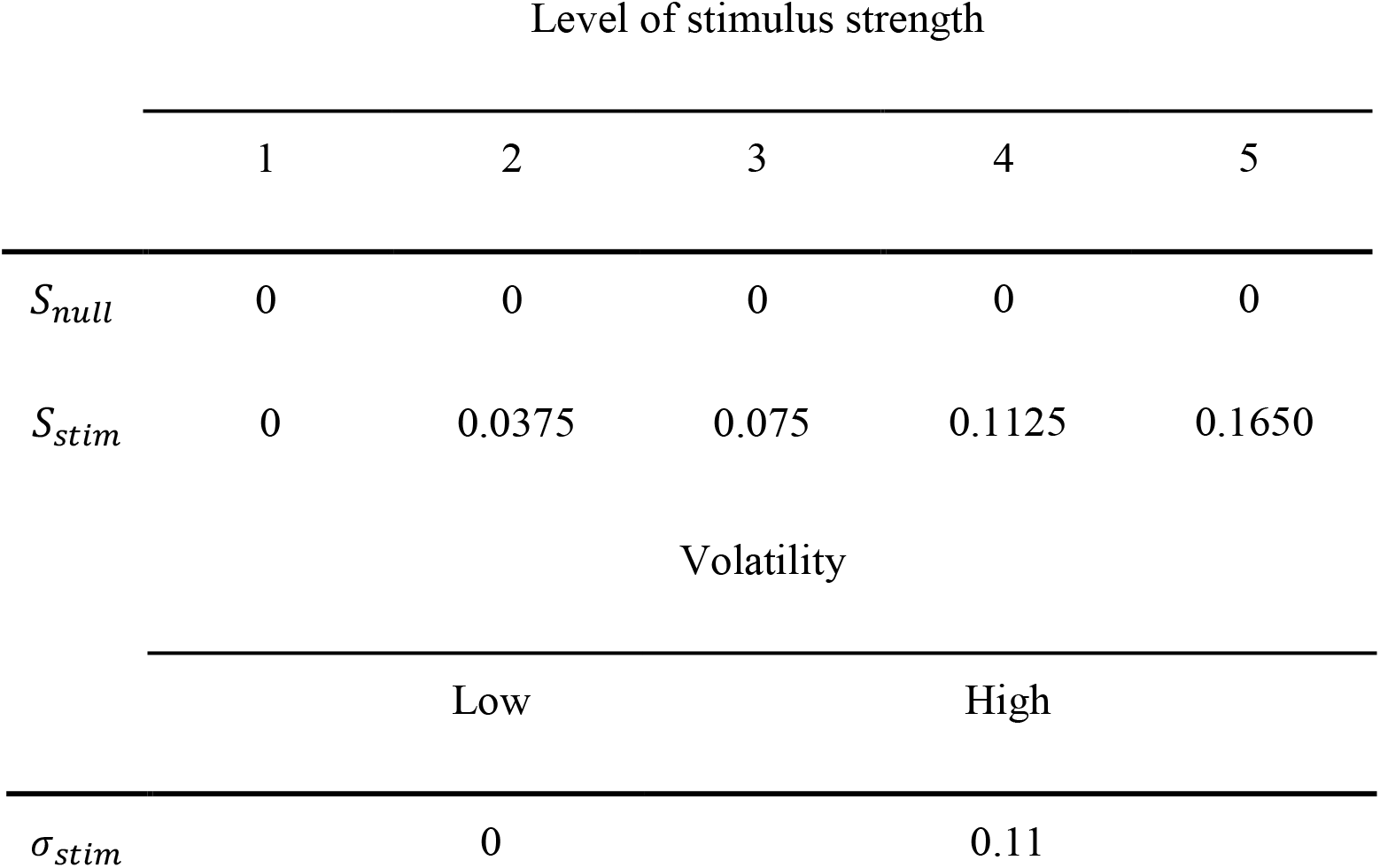
Simulation-specific parameter values used for simulating the results of (Zylberberg et al., 2016).

To convert the model’s continuous confidence variable C to confidence ratings, we set N_ratings_ = 2 to match the confidence scale used in Zylberberg et al. Because it was not possible to achieve a close numerical fit to the empirical data, we arbitrarily set *U_r_* to the median of the confidence distribution of C, and similarly for C*.

### 3. Rhesus macaque electrophysiology

#### 3.1. Behavioral task

The task was a classic dot-motion discrimination task. One monkey observed random dot motion stimuli and made saccades to targets on the screen to indicate its choice about the dot motion direction. On each trial, after an initial fixation period two choice targets appeared for a random time around 500 ms (drawn from an exponential distribution to avoid prediction, Figure 5a). A random dot motion stimulus subsequently appeared at the center of the screen with some percentage of dots moving in the same direction; this is the ‘coherence’ of the dot motion. After the dot motion stimulus offset, the monkey made a saccade either towards the Target (in the recorded neurons’ response field) or towards a Distractor (opposing the response field).

**Figure 5.**
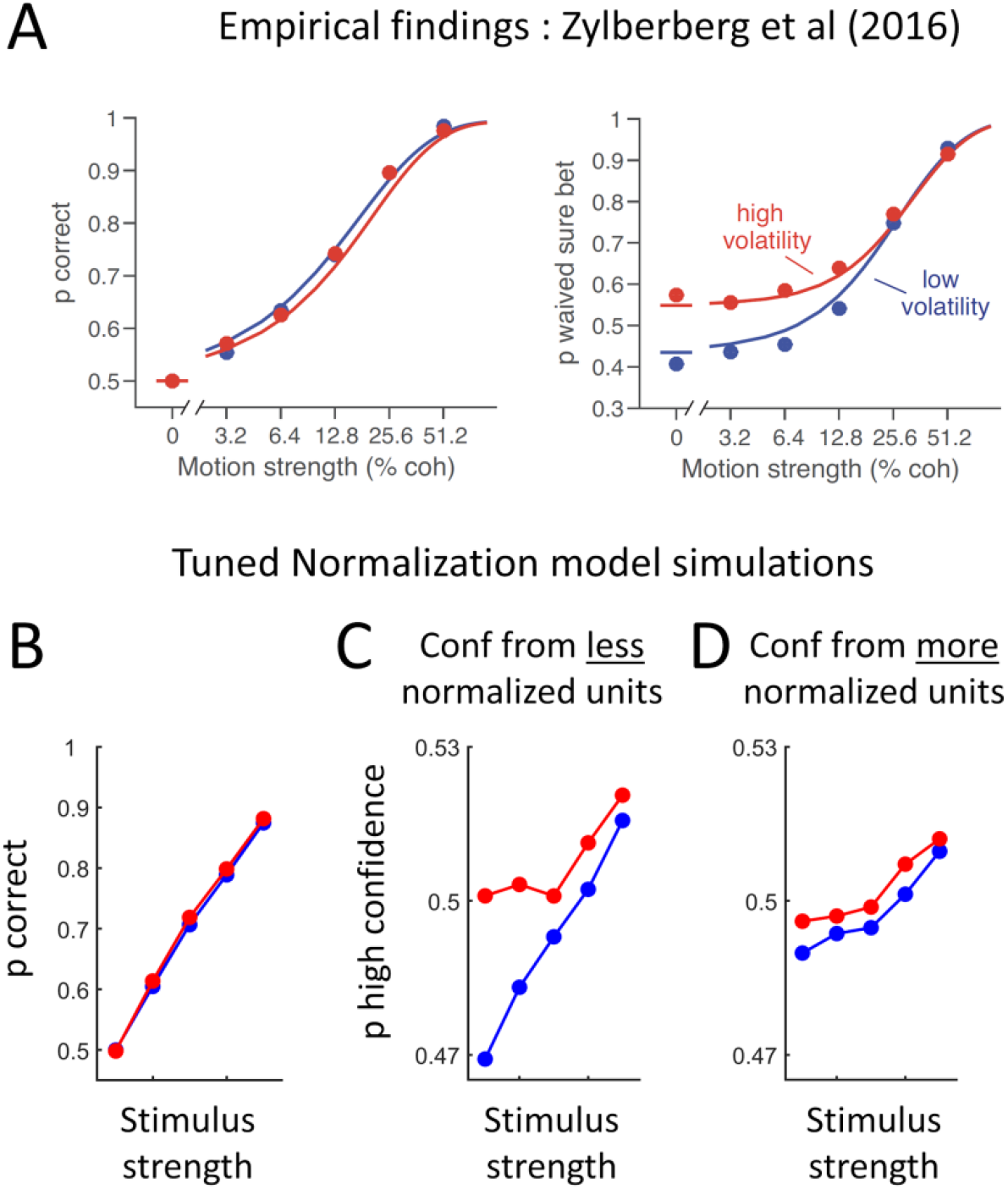
The Tuned Normalization model predicts the effect reported by Zylberberg and colleagues (Zylberberg et al., 2016). (a) The authors showed that increased stimulus volatility leads to similar objective performance but increased confidence ratings, especially at low objective performance levels. This figure is reproduced from (Zylberberg et al., 2016). (b, c) The Tuned Normalization model qualitatively reproduces these effects. (d) The effect of stimulus volatility on confidence was nearly abolished in control simulations in which more normalized units are weighted more heavily for computing confidence, suggesting that the result in (b) critically depends on confidence being computed primarily from less normalized units. Cohen’s d for confidence effects in the main model simulations was ~1.5 – 6 times greater than in the control simulations across all levels of stimulus strength.

Head restrained monkeys (see below) sat in a custom-sized chair facing a CRT monitor (1024 × 768 pixel resolution, 85 Hz refresh rate) at a distance of 37 cm. A photocell secured to the monitor sent a transistor-transistor logic pulse to the PC used to display stimuli to provide an accurate measure of stimulus event timing. Each trial began by the monkey looking at a white dot that appeared at the center of the monitor. After a brief delay of ~500 ms, randomized from an exponential distribution to prevent prediction, a red target appeared in one hemifield, and a green target appeared in the other. Targets were isoluminant (13 cd/m^2^), and the hemifield in which the red or the green choice target appeared varied randomly from trial-to-trial (Ferrera et al., 2009; Bennur and Gold, 2011). After a second randomized delay (600-1050 ms), a random dot motion kinematogram stimulus (pattern diameter = 60, 26 cd/m^2^; dot size=0.10; dot separation=0.1820; total density=5 dots/deg^2^) appeared at the center of the monitor together with the white fixation point and remained on the screen for 200 ms. The stimulus’s disappearance was followed by a 500-600 ms delay (the exact value was drawn from an exponential distribution to avoid prediction), at which point the fixation point also disappeared, instructing the monkey to report its choice with an eye movement.

Although there were other trial types presented for the purposes of the experiment for which the data were originally collected, for our analysis we focused on the ‘catch trials’ in which 100% of the dots were moving in the preferred direction (i.e., toward the Target, ‘100% Preferred’ trials) or 50% toward the Target and 50% toward the Distractor (‘50-50% Preferred + Non-Preferred’ trials). These 100% preferred and 50-50% motion ‘catch trials’ were sparsely interspersed among other trial types in the main task, so the monkey completed 154 100% ‘Preferred’ trials and 149 50-50% ‘Preferred + Non-Preferred’ trials in total across eight days. However, because the electrode was placed in a different location on each recording day, on average only 19.25 100% ‘Preferred’ and 18.63 50-50% ‘Preferred + Non-Preferred’ trials were collected for each unit (Table 5). Unfortunately, electrode placement techniques also precluded precise neuroanatomical localization of the electrode -- and thus the recorded neurons -- on each recording day.

**Table 5.** Daily specifics for recordings from macaque superior colliculus (SC): number of SC neurons recorded after spike-sorting (see Methods, main text), and number of trials of ‘Preferred’ and ‘Preferred + Non-Preferred’ stimuli, respectively.

| **Day** | **# units** | **# ‘Preferred’ trials** | **# ‘Preferred + Non-Preferred’ trials** |
| --- | --- | --- | --- |
| **1** | 31 | 15 | 13 |
| **2** | 24 | 19 | 16 |
| **3** | 16 | 20 | 20 |
| **4** | 15 | 20 | 20 |
| **5** | 18 | 20 | 20 |
| **6** | 17 | 20 | 20 |
| **7** | 19 | 20 | 20 |
| **8** | 16 | 20 | 20 |

#### 3.2. Electrophysiological recordings

##### Surgical procedures

One male rhesus monkey weighing approximately 10 kg was prepared for electrophysiological recordings and measurement of eye movements. A headpost was implanted to secure the head, and an MRI-compatible recording chamber (Crist instruments, MD) was placed at AP +3, ML 0 and angled 38° posteriorly to access the superior colliculus (SC). Precise positioning of the headpost and the recording chamber was obtained using MRI-guided surgical software (BrainSight, Rogue Research, Montreal, CA). To track eye movements, eye position was monitored with an iView camera (Sensomotoric instruments, Boston, MA). All surgical procedures were performed under general anesthesia using aseptic procedures. Anesthesia was induced with ketamine and midazolam (5.0 mg/kg and 0.2 mg/kg, i.m.). Atropine (0.04 mg/kg, i.m.) was provided to reduce salivation. The monkey was intubated and maintained under general anesthesia with isoflurane. One hour before the procedure, the animal received buprenorphine (0.01 mg/kg, i.m) and the antibiotic Excede (20 mg/kg, i.m; 7 days slow release) and then meloxicam (0.3 mg/kg, i.m) at the conclusion of the procedure. Meloxicam (0.2 mg/kg, i.m) and buprenorphine (0.01mg/kg, i.m) were administered for 3 days post-surgically for multimodal analgesia. All experimental protocols were approved by the UCLA Chancellor’s Animal Research Committee (IACUC) and complied with and generally exceeded standards set by the Public Health Service policy on the humane care and use of laboratory animals.

##### Eye movement recordings

Experiments used a QNX-based real-time experimental data acquisition and visual stimulus generation system, Rex and Vex, developed and distributed by the Laboratory of Sensorimotor Research National Eye Institute in Bethesda MD (Hays et al., 1982) to create the behavioral paradigm and acquire eye position data. The camera acquired eye position signals were filtered digitally using a built-in bilateral filter. We used an automated procedure to define the onset of saccadic eye movements using eye velocity (20°/s) and acceleration criteria (5000°/s2), respectively. The adequacy of the algorithm was verified and adjusted as necessary on a trial-by-trial basis by the experimenter.

##### Neuronal recordings

We recorded single neurons and multineuron activity in the SC with a 16 channel platinum/iridium V Probe coated with polyimide (Plexon, Dallas, TX), with an impedance of 275 (±50) kΩ. The V Probe was inserted through a guide tube, perpendicular to the surface of the SC, positioned with a grid system (Crist et al., 1988) and advanced using an electronic microdrive system controlled by a graphical user interface (Nan Instruments, Israel). Action potential waveforms were bandpass filtered (250 Hz – 5 kHz; 4 pole Butterworth), and amplified using the BlackRock NSP hardware system controlled by the Cerebus software suite (BlackRock Microsystems, Utah). Neurons were isolated online using time and amplitude windowing criteria. The times of occurrence of action potentials were digitized at 16 bit resolution and sampled at 1 kHz and saved to disk. Neuronal waveform data were digitized at 16 bit resolution and sampled at 30 kHz and saved to disk for offline analysis using Plexon offline sorting algorithms (Plexon, Dallas, TX).

Response fields (RF) of SC neurons were mapped online. Mapping was done by moving a spot around the monitor and having monkeys make delayed saccades to the different spots. During this mapping procedure, on each trial a fixation spot appeared initially at the center of the screen, and monkeys fixated for 500-1000 ms. A second spot then appeared peripherally while monkeys remained fixated for another 200-400 ms until the fixation spot disappeared; the location of this dot was controlled by mouse movements, and could be anywhere on the screen. Delay times were randomized, drawn from an exponential distribution to prevent prediction. The fixation spot’s disappearance cued the monkey to make a saccade to the peripheral target. If he made a correct saccade (within a window of 2° diameter), he received a fluid reward of 0.1mls.

We listened for maximal discharge for each saccade on-line. The center of the RF was considered to be the location at which a saccade was associated with maximal audible discharge of the neuron. The center of the RF was confirmed by plotting the discharge rate as a heat map in Cartesian coordinates. Only neurons with RF eccentricities between 7 and 20° were selected for further study in order to ensure no overlap of the RF with the centrally-located motion cue stimulus. Although electrode penetration was aimed at the SC perpendicular to its surface, we noticed slight differences in the RF of each recording site of the V-Probe, during the same penetration. Therefore, the RF was optimized for at least one recording site on each experimental day.

#### 3.3. Calculating normalization tuning via the Modulation Index

Drawing from previous work (Ni et al., 2012; Ruff et al., 2016; Verhoef and Maunsell, 2017), we calculated the normalization tuning Modulation Index (MI) as:

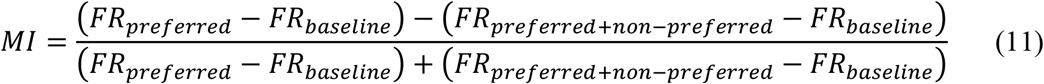

For the baseline firing rate, we took advantage of the fact that the conditions of interest were interspersed in a larger dataset, such that there were many other trials we could use to calculate baseline. Therefore, we calculated each neuron’s baseline firing rate as the average firing rate across all trials (including those in the unrelated task) in the 400 ms prior to the motion cue onset. We calculated the firing rate during the presentation of ‘Preferred’ or ‘Preferred + NonPreferred’ motion as as the average firing rate across trials during the first 400 ms after motion onset in the relevant trial types. We acknowledge that this definition of Modulation Index in some ways departs from that used in some previous reports to identify normalization tunedness (Ni et al., 2012; Ruff et al., 2016; Verhoef and Maunsell, 2017); however, the small number of relevant trials in this existing dataset unfortunately precluded adoption of previously-used metrics.

Action potential waveforms were sorted offline using the Plexon Offline Sorter (Offline Sorter, Plexon Inc). In total, usable signal was recorded from 156 neurons. Because we were interested in the degree to which a neuron modulates its firing rate in the presence of ‘Preferred’ versus ‘Preferred + Non-Preferred’ motion stimuli, we excluded neurons that demonstrated no significant response to dot motion stimulus onset, defined as a firing rate during stimulus presentation equal to or less than the baseline firing rate. This left 125 neurons for the calculation of MI.

We calculated MI for all trials, and also split trials randomly by whether the trial was an even- or odd-numbered trial and calculated MI separately for these subsets of trials. To check for the presence of normalization tuning, we correlated the MI calculated for even versus odd numbered trials across all neurons.

#### 4.3. Code and Data Accessibility

Code for the Tuned Normalization model simulations, as well as data and code for the monkey electrophysiology, will be shared upon reasonable request.

## Results

### The Differential Tuned Normalization model accounts for behavioral findings

We examined model predictions for three different behavioral effects reported in the literature by simulating the tasks described in those papers with the Differential Tuned Normalization model. For each report from the literature, we selected inputs that would mimic the stimulus seen by human or monkey observers, and ran a simulated ‘observer’ through 100,000 simulated trials per condition.

Critically, we fixed all internal parameters of the model for all simulations, such that the network architecture was identical for all tasks and only the inputs (i.e., ‘stimuli’), and confidence thresholds tuned to those inputs, changed from one simulation to the next. Confidence thresholds were chosen for each simulation to approximate the fitted data. For a given simulation and choice of internal confidence variable C (Equations 7 and 9), the same confidence thresholds were applied to simulated data from all stimulus conditions, so that any between-condition differences in confidence arose purely from the influence of network activity on C. Changing confidence thresholds mainly affects the mean confidence values, while retaining similar standard deviations and between-condition differences. Therefore, the effect size of differences in confidence between conditions was robust to choice of confidence thresholds.

As discussed above, our central hypothesis is that decision-congruent evidence effects in confidence rating could arise as a result of less normalized units being weighted more heavily in the computation of confidence than in the computation of perceptual decisions. To assess this hypothesis, we conducted control simulations in which the confidence decision variable was identical to the decision variable used for stimulus discrimination, i.e. one in which *more* normalized units rather than *less* normalized units were weighted most heavily. If our hypothesis is correct, then we would expect to find that decision-congruent confidence effects are more pronounced when confidence is rated using less normalized units.

To anticipate, we found that when confidence more heavily depended on *less* normalized units, our model simulations could reproduce decision-congruent confidence effects found in the literature. By contrast, when confidence more heavily depended on *more* normalized units, decision-congruent confidence effects were almost completely abolished. These findings lend support to the hypothesis that differential contributions of differently normalized neurons to stimulus discrimination and confidence can provide a neural mechanism for decision-congruent confidence effects.

#### (1) Controlling positive and negative evidence to yield equal performance and unequal confidence

Koizumi and colleagues (Koizumi et al., 2015) reported that certain stimulus conditions can produce the same decisional accuracy but different confidence. Human observers viewed compound stimuli consisting of two superimposed oriented sinusoidal grating targets embedded in noise, with one grating (“positive” evidence) at higher contrast than the other (“negative” evidence) (Figure 3a). Observers indicated whether each stimulus’ primary orientation was tilted left or right of vertical and provided a confidence judgment. Four conditions were presented in a 2×2 design: High or Low Positive Evidence, and Difficult or Easy. Orientation discrimination task performance was equivalent across the “High Positive Evidence” and “Low Positive Evidence” conditions within a difficulty level, but the “High Positive Evidence” condition led to higher confidence judgments (Figure 3b).

We simulated two Difficult and two Easy conditions, crossed with Low versus High Positive Evidence (Table 2). We treated stimulus strength for positive evidence in the four conditions as free parameters. Following the methods in Koizumi et al., we constrained stimulus strength for negative evidence to be NE = 0.7*PE in the High Positive Evidence conditions and NE = 0.35*PE in the Low Positive Evidence conditions. We found that the Tuned Normalization model reproduced all aspects of the behavior reported by Koizumi and colleagues (Koizumi et al., 2015): easier conditions led to higher confidence, and High Positive Evidence led to higher confidence than Low Positive Evidence (Figure 3c).

In the control simulation where more heavily normalized units determined confidence, the effect of High vs Low Positive Evidence on confidence was nearly abolished (Figure 3d). We quantified this by computing Cohen’s d for confidence in the High vs Low Positive Evidence conditions separately for the Difficult and Easy conditions. The main model simulations yielded Cohen’s d values of 0.29 and 0.40 in the Difficult and Easy conditions, whereas these values decreased to near-zero values of 0.06 and 0.08 in the control simulations.

Interestingly, despite the highly constrained nature of the model parameter specifications, including the *a priori* specification of the ratio of stimulus strength for positive and negative evidence values, the Cohen’s d for confidence effects in the simulations was similar to the average within-subject^1^ Cohen’s d for confidence effects in the empirical data (0.33 and 0.26 in the Difficult and Easy conditions, respectively, as computed from raw data provided by the authors; cf. simulated values of 0.29 and 0.40, respectively).

#### (2) Introducing noise: TMS, inattention, or microstimulation

It has been reported that TMS to visual cortex reduces human observers’ ability to perform an objective discrimination task, but increases their confidence/visibility judgments (Rahnev et al., 2012b; Peters et al., 2017a). Another similar approach used attention to manipulate the precision of the perceptual signal (Rahnev et al., 2011). After matching discrimination performance for attended versus unattended Gabor patch stimuli embedded in noise, the authors found that observers tended to report higher visibility for the unattended (i.e., noisier) stimuli. These effects also appear similar to the recent report that microstimulation to neurons can increase confidence while slightly reducing objective task performance (Fetsch et al., 2014). Rhesus macaques performed a dot-motion discrimination task in which some trials provided an opt-out sure-reward option; it is assumed monkeys selected this opt-out option when they were unsure of the direction of the dot motion (Kiani and Shadlen, 2009). On some trials, microstimulation was applied to neurons in area MT/MST. Microstimulation slightly reduced task performance, but caused monkeys to opt out less often, meaning they had higher confidence.

We hypothesized that these effects all depend on a similar computational mechanism, and that the Tuned Normalization model would likely be able to predict them all. To test this, we simulated a discrimination task in which stimulus strength remained constant across two conditions, but one condition had increased levels of additive noise in evidence accumulation (Table 3), mimicking the effects of TMS (Rahnev et al., 2012b; Peters et al., 2017a), inattention (Rahnev et al., 2011), and microstimulation (Fetsch et al., 2014). We present the results in comparison to figures reproduced from Rahnev et al. (Rahnev et al., 2012b), since they provide a succinct summary of this behavioral effect (Figure 4a). For the Tuned Normalization model, increased evidence accumulation noise led to lower objective discrimination performance (Figure 4b) but higher confidence (Figure 4c), mimicking empirical effects on performance and confidence induced by TMS, inattention, and microstimulation. Importantly, the choice of the single parameter (evidence accumulation noise σadd) selected to yield closest match to the empirical difference in d’ between the “noisier” (i.e., TMS, inattention, or microstimulation) versus “less noisy” conditions simultaneously yielded a close match to the empirical difference in confidence without any further intervention in model parameters. That is, we did not fit this *σadd* parameter separately to the objective d’ and confidence results; rather, the confidence results follow directly from selecting the *σadd* magnitude that best fits the objective d’ behavior.

In the control simulation where more heavily normalized units determined confidence, the effect of added evidence accumulation noise on confidence was nearly abolished (Figure 4d). We quantified this by computing Cohen’s d for confidence in the Added Noise vs Baseline conditions (corresponding to the Occipital TMS vs Vertex TMS conditions in Rahnev et al. 2012). The main model simulation yielded Cohen’s d = 0.1, whereas this value decreased by half to 0.05 in the control simulation. We conducted additional simulations to verify that Cohen’s d was robustly larger in the main simulation than in the control. In the simulation results reported above, the additive stimulus noise parameter of the model, σadd, was increased from 1 in the Baseline condition to 1.05 in the Added Noise condition. In five supplementary simulations, we set *σadd* to values of 1.1, 1.15, 1.2, 1.25, and 1.3 and recorded the Cohen’s d in the main and control simulations. Across-condition differences in simulated d’ and confidence increased with increasing *σ_add_*. Cohen’s d reached a maximum value of 0.56 at the maximum tested value of *σ_add_* = 1.3, and was twice as large as Cohen’s d in each control simulation (regression slope = 2.02). Thus, for a given level of difference in task performance, the effect size of the difference in confidence was robustly larger when confidence was computed primarily from less rather than more heavily normalized units.

#### (3) Stimulus volatility leads to equal performance but increasing confidence

Zylberberg and colleagues (Zylberberg et al., 2016) reported that introducing stimulus *volatility* produced negligible decrement in objective performance capacity, but an overall increase in confidence in the more volatile conditions. To introduce stimulus volatility, they used a random dot motion stimulus in which the coherence of the dot motion was held constant in the Low Volatility condition, whereas in the High Volatility condition this coherence represented the mean of a distribution of possible coherences that could change on a frame-by-frame basis. Macaques and human observers judged the primary direction of the dot motion and either rated confidence (humans) or used an opt-out sure-reward target to indicate confidence (monkeys). Higher stimulus volatility led to only a small decrease in objective performance but an increase in confidence, particularly at lower levels of stimulus strength (Figure 5a). We simulated their task by introducing variability in stimulus strength in each time step of the evidence accumulation process (Table 4), and found that the Tuned Normalization model was able to capture the qualitative patterns in Zylberberg et al.’s findings (Figure 5b, c).

In the control simulation where more heavily normalized units determined confidence, the effect of added stimulus volatility on confidence was nearly abolished (Figure 5d). We quantified this by computing Cohen’s d for confidence in the High Volatility vs Low Volatility conditions. The main model simulation yielded Cohen’s d = 0.064, 0.040, 0.016, 0.018, and 0.010 across the five levels of stimulus strength. These values decreased to 0.012, 0.007, 0.008, 0.012, and 0.005 in the control simulations. Thus, at the two lowest levels of stimulus strength in which confidence effects due to volatility in the main model were most apparent, the effect size of volatility was over five times greater than in the control simulations. At the higher stimulus strength levels, confidence effects remained about two times greater in the main model simulations than in the control simulations.

### Tuned normalization in a decision-making area: the superior colliculus

Based on the results of our simulations, one may expect that neurons in decision-making areas show this pattern of tuned normalization. Although tuned normalization has been observed in primary sensory cortical areas, including V1, V4, and MT/MST (Ni et al., 2012; Ruff et al., 2016; Verhoef and Maunsell, 2017), until now it has not been observed in cortical areas known to contain neurons that accumulate evidence (e.g., LIP, FEF) or any subcortical structures, including those with similar function (e.g., superior colliculus; (Gold and Shadlen, 2000; Horwitz et al., 2004; Smith and Ratcliff, 2004; Kim and Basso, 2008)).

We capitalized on an existing dataset to look for tuned normalization in Rhesus macaque superior colliculus (SC). In a task designed for a different study, one Rhesus macaque engaged in a dot motion direction discrimination task (Figure 6a) while data were recorded from 156 SC neurons using a multi-channel V-probe. The response fields of the SC neurons had previously been mapped, and saccade targets that the monkey could use to indicate its dot motion direction choice were placed either in each neuron’s response field (‘Target’) or in a location on the opposite side of the screen (‘Distractor’). On each trial, a Target and Distractor appeared on the screen followed by centrally-presented dot motion, and the monkey made saccades to the Target or Distractor to indicate its choice (Figure 6a).

**Figure 6.**
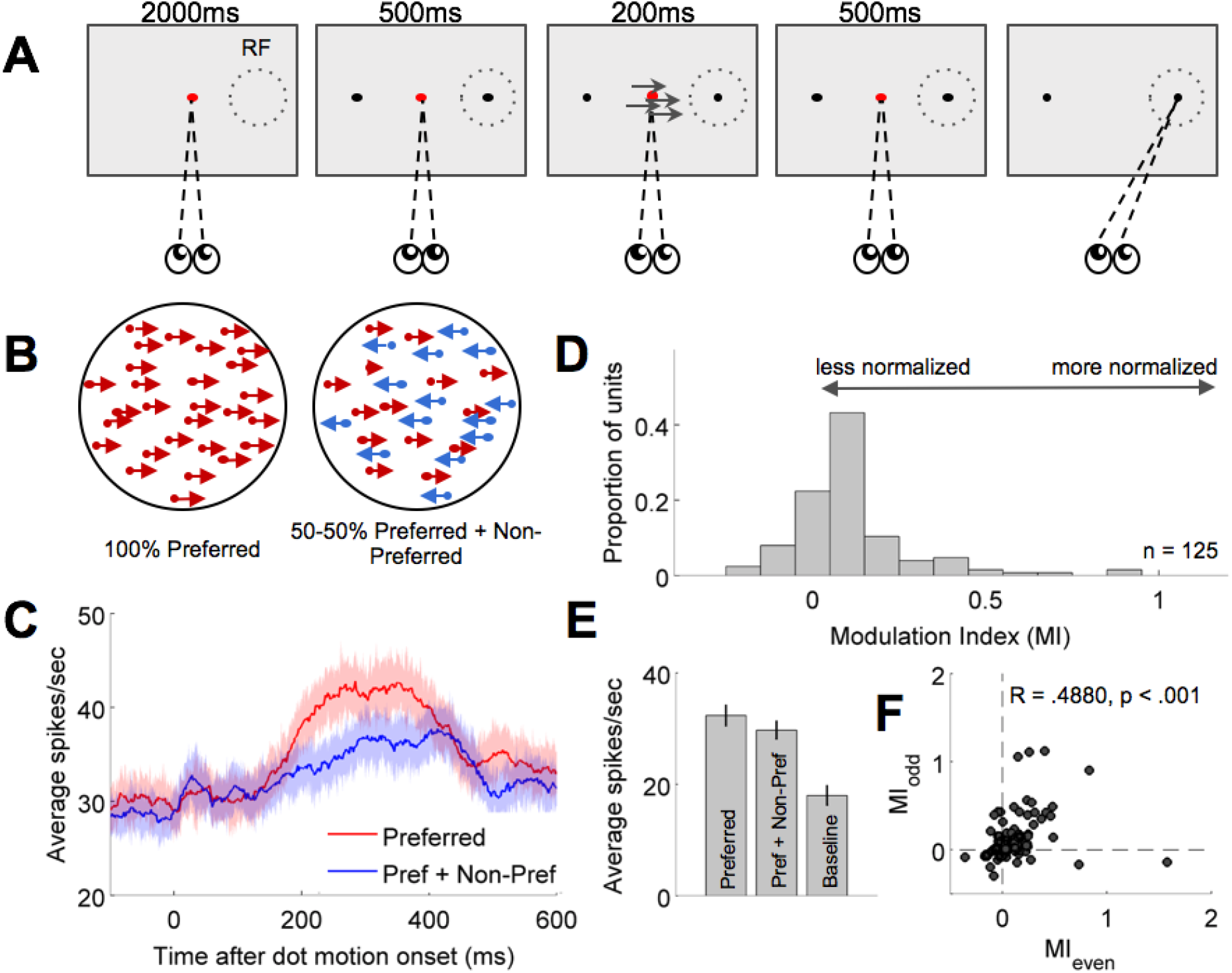
Tuned normalization can be observed in macaque superior colliculus (SC). (a) We took advantage of an existing dataset in which one monkey performed a dot motion discrimination task. The monkey’s SC neurons’ response fields (RF) were mapped prior to the beginning of the task on each day. On each trial, following a fixation period a brief dot motion stimulus appeared. The monkey observed the stimulus, and then after a delay period made a saccade to the target indicating its dot motion direction discrimination decision. (b) To look for tuned normalization, we focused on ‘catch trials’ in which 100% ‘Preferred’ motion (indicating a saccade towards the neuron’s RF) or 50-50% ‘Preferred + Non-Preferred’ motion was presented to the monkey. These trials allowed us to calculate each neuron’s Modulation Index (MI; see Online Methods, Section S2.3) to measure each neuron’s degree of tuned normalization. (c,e) As expected, neurons showed higher firing rates to ‘Preferred’ stimuli than to mixed ‘Preferred + NonPreferred’ stimuli in the post-stimulus window. These were also higher than the baseline (prestimulus) firing rate. Mean firing rates in (e) were computed from the 400 ms before (baseline) and after (Preferred and Preferred + Non-Preferred) stimulus onset. (d) We observed a wide distribution of MI values for the SC neurons we recorded, with a median significantly greater than 0 (Wilcoxon sign-rank test: z = 6.9224, p < .001). (f) To confirm that the observed distribution of MI reflected tuned normalization and not random variation, we split the trials randomly into those with even versus odd trial numbers, and re-calculated MI for each neuron. These disjoint datasets produced MI values that were strongly correlated across even versus odd trials (R = .4880, p < .001), indicating that the degree to which a neuron is normalized is both *different* across neurons and *consistent* within individual neurons. These results provide evidence of tuned normalization outside the cortex, and that it exists in evidence-accumulation neurons as opposed to just in primary sensory neurons as previously observed (Ni et al., 2012; Ruff et al., 2016; Verhoef and Maunsell, 2017).

‘Catch trials’ of 100% coherent dot motion in the ‘Preferred’ direction (i.e., the direction of motion that should lead to a saccade towards the Target) of each SC neuron, and other catch trials consisting of transparent motion that was 50% ‘Preferred’ and 50% ‘Non-Preferred’ (i.e., the direction of motion that should lead to a saccade towards the Distractor) were interspersed among the trials for that separate study (Figure 6b). We measured the firing rate responses of SC neurons in response to the ‘Preferred’ and 50-50% ‘Preferred + Non-Preferred’ dot motion stimuli in these catch trials, as well as their baseline firing rates, and focused on neurons that exhibited evidence accumulation.

As expected, neurons showed stronger and earlier spiking for ‘Preferred’ stimuli than for ‘Preferred + Non-Preferred’ (Figure 6c). The average of both of these firing rates in the first 400 ms after stimulus onset were above baseline firing rates (Figure 6e).

Importantly, these trial types can be used to examine normalization tuning via calculation of a ‘Modulation Index’ (MI) for each recorded neuron (Equation 11). Following previous convention (Ni et al., 2012; Ruff et al., 2016; Verhoef and Maunsell, 2017), the MI quantifies the degree to which an accumulation neuron with a given tuning preference (i.e., response field) exhibits modulation in its firing rate due to lateral inhibition when presented with simultaneous ‘Preferred + Non-Preferred’ versus ‘Preferred’ stimuli alone. Smaller MI means a cell is less normalized, because the neuron’s response to Preferred stimuli is relatively independent of the presence of Non-Preferred stimuli; in contrast, larger MI indicates a cell is more normalized, because the neuron’s firing rate is strongly reduced in the presence of simultaneous Preferred and Non-Preferred motion over its firing rate to Preferred motion alone.

125 neurons demonstrated meaningful responses to dot motion stimulus presentation and evidence accumulation properties (Figure 6c; see also Materials & Methods, Section 3). The distribution of MI for these neurons was centered above zero (μ = 0.1063, σ = 0.1747) (Figure 6d). Because a Lilliefors test revealed that MI significantly deviated from a normal distribution, we used the Wilcoxon signed-rank test (Wilcoxon, 1945) to demonstrate that this distribution is centered significantly higher than zero (z = 6.9224, p < .001). This indicates that SC neurons exhibit divisive normalization as expected, congruent with previous reports (Schiller and Koerner, 1971; Sterling, 1971; Goldberg and Wurtz, 1972; Moors and Vendrik, 1979; Basso and Wurtz, 1998; Li and Basso, 2005; Phongphanphanee et al., 2014; Vokoun et al., 2014).

To determine the extent to which fluctuations in MI reflected tuned normalization (i.e., different and *consistent* degrees of normalization tuning for each individual neuron) and not just random noise, we split trials into even versus odd trial numbers and calculated MI for each disjoint subset of trials. We then examined the correlation between even and odd trials as a robust way to identify tuned normalization as distinct from noise. Importantly, this revealed a highly significant correlation between even and odd trials (R = 0.4880, p < .001), indicating that the tuned normalization exhibited by each neuron is highly consistent across trials (Figure 6f).

These results demonstrate that tuned normalization exists in evidence-accumulation neurons in decision-making circuits, suggesting that the Tuned Normalization model framework proposed here may indeed be capable of implementing the calculation of subjective confidence. We do acknowledge that these preliminary results depend on relatively few trials (in an existing dataset) and in some ways depart from other methods by which tuned normalization has been identified in previous reports (Ni et al., 2012; Ruff et al., 2016; Verhoef and Maunsell, 2017); further, we also note that the observation of tuned normalization in accumulation neurons does not necessitate that such neurons *must* encode decisions versus confidence in the manner hypothesized here. However, these results provide an initial proof of concept, paving the way for future electrophysiological studies examining the predictions of our model in areas previously related to perceptual confidence, such as LIP (Kiani and Shadlen, 2009). Our findings also provide the first demonstration of normalization tuning (as opposed to just the presence of normalization) in a subcortical structure, as previously it has been observed only in cortical areas (Ni et al., 2012; Ruff et al., 2016; Verhoef and Maunsell, 2017).

## Discussion

How the brain calculates subjective decision confidence is still a topic of active debate (Ma et al., 2006; Fleming and Dolan, 2012; Yeung and Summerfield, 2012; Charles et al., 2013; Orbán et al., 2016; Pouget et al., 2016; Sanders et al., 2016). Although dominant models suggest that confidence reflects an optimal readout of the probability that a decision is correct (Ratcliff and Rouder, 1998; Ratcliff and McKoon, 2008; Pleskac and Busemeyer, 2010; Tsetsos et al., 2012; Fetsch et al., 2014; Kiani et al., 2014; Pouget et al., 2016; Sanders et al., 2016; Zylberberg et al., 2016), it appears challenging for such models to account for counterintuitive behaviors in which confidence and accuracy do not covary (Rahnev et al., 2011, 2012a, 2012b; Koizumi et al., 2015; Maniscalco et al., 2016; Samaha et al., 2016). An alternative hypothesis suggesting that confidence reflects a heuristic reliance on decision-congruent evidence (Zylberberg et al., 2012; Koizumi et al., 2015; Maniscalco et al., 2016; Samaha et al., 2016, 2017) captures many of these behaviors, and is supported by human intracranial electrophysiology (Peters et al., 2017b).

Here, we considered how decision-congruent computations of perceptual confidence might be biologically implemented based on known properties of perceptual circuitry. We hypothesized that tuned normalization (Ni et al., 2012; Ruff et al., 2016; Verhoef and Maunsell, 2017) differentially influences a neuron’s role in perceptual decision-making and confidence, such that more normalized units (corresponding to the net evidence for a perceptual choice) drive decisions and less normalized units (corresponding to decision-congruent evidence) drive confidence. We developed the Differential Tuned Normalization model to test this hypothesis. Our results show that such a network can explain counterintuitive behaviors reported in the literature (Rahnev et al., 2011, 2012a; Fetsch et al., 2014; Koizumi et al., 2015; Samaha et al., 2016; Zylberberg et al., 2016). We further demonstrate that the model’s special property of weighting less normalized units more heavily in computing confidence is the key to capturing empirical findings, since control simulations demonstrate that the model fails to reproduce these findings when instead *more* normalized units drive confidence. Finally, we used electrophysiological recordings in monkeys to reveal this neuronal property in an area containing the type of evidence accumulation neurons typically assumed to be involved in perceptual decision-making (Ratcliff and Rouder, 1998; Usher and McClelland, 2001; Ratcliff and McKoon, 2008; Pleskac and Busemeyer, 2010; Yeung and Summerfield, 2012, 2014; Kiani et al., 2014). This provides preliminary but converging evidence that decision-congruent confidence computations may be implemented via tuned normalization.

It might be argued that some over-simplified optimal diffusion-type models (Ratcliff and Rouder, 1998; Ratcliff and McKoon, 2008; Pleskac and Busemeyer, 2010) should not be expected to account for counterintuitive behaviors due to their simplicity. A modification of these optimal diffusion-type models was recently proposed, suggesting that the optimal confidence readout must also depend on the time it took for evidence to accumulate (Fetsch et al., 2014; Kiani et al., 2014; Zylberberg et al., 2016). Although it has been suggested that neurons in lateral intraparietal cortex (LIP) may encode elapsed time (Leon and Shadlen, 2003; Janssen and Shadlen, 2005; Finnerty et al., 2015), these neurons’ activity has not yet been causally or directly linked to subjective confidence (see also (Bang and Fleming, 2018)). This suggests that how this time-dependent diffusion framework might be biologically implemented is nontrivial, inspiring the work presented here.

It should be noted that we did not perform exhaustive fitting of model parameters to each individual data set. Instead, we took the simpler approach of fixing most model parameters across all simulations (see Materials and Methods, Section 1; Table 1). The only parameters free to vary across simulations were (1) within-condition stimulus strength (to fit task performance), (2) within-condition noise in neural activity (simulation 2) or stimulus strength (simulation 3) (to fit across-condition differences in task performance and confidence), and (3) confidence thresholds constrained to be identical across all conditions within a simulation (to scale overall confidence values) (see Materials and Methods, Section 2; Tables 2-5).

Importantly, these modeling decisions revealed that the Tuned Normalization model can reproduce the quantitative and qualitative effects reported in the literature with a *single* set of parameters controlling network dynamics: only the task-specific inputs changed from one simulation to the next (mimicking the different stimuli and confidence scales in each behavioral task), while the internal network architecture remained identical across simulations. Our simplified approach to model fitting had the further virtue of converting a potentially underconstrained model fitting project into a highly constrained one that mitigated against overfitting and facilitated comparison of results across simulations. Our goal with this model, therefore, was to provide a proof of concept that tuned normalization in sensory circuits can provide a biologically plausible mechanism for implementing decision-congruent confidence computations. It is therefore somewhat remarkable that, in spite of these parameter fitting constraints, the model was able to closely capture exact empirical quantities in two simulations (Figures 3 and 4) as well as the qualitative pattern of data in a third simulation (Figure 5).

Importantly, our results suggest a potential adaptive purpose for tuned normalization (Ni et al., 2012; Ruff et al., 2016; Verhoef and Maunsell, 2017) within a behaving organism. It appears that the presence of both more and less normalized neurons within a perceptual decision-making circuit may allow an organism to better solve both fine-grained discrimination and detection tasks. When making fine-grained discrimination or identification judgments about an object or stimulus (“What is that thing?”), an organism should rely on a system that is not as sensitive to random fluctuations, i.e. a more strongly normalized system. But when making detection decisions (“Is there something out there?”), such strong lateral inhibition would be highly undesirable, so an organism should rely on less normalized parts of the network. Both of these tasks are important for an organism to execute, and so it seems unlikely that a system would only be optimized for one or the other.

The question then becomes why the system would opt to recruit the ‘detection’ portions of its circuitry to compute confidence, specifically relying on the magnitude of decision-congruent evidence. One reason may simply be heuristic, that the detectability and identifiability of a stimulus are almost always perfectly correlated in the real world: perfectly detectable but not discriminable (or vice versa) stimuli are rare outside the laboratory. Indeed, it has been noted that the width of the posterior distribution in a probabilistic population code (Ma et al., 2006, 2008) covaries with the overall firing rate of a population (Bays, 2016); less normalized ‘detection’ neurons would more readily affect a population’s overall firing rate, suggesting a potential neural substrate for this heuristic. Perhaps, in the interest of computational efficiency, the system has come to rely on this heuristic, which minimizes the need to retain information about unchosen stimulus identity possibilities once a perceptual inference has been made (Peters et al., 2017b). Indeed, such over-reliance on decision-congruent evidence -- i.e., a “confirmation bias” (Abrahamyan et al., 2016) -- has also been observed in other post-decisional (non-metacognitive) perceptual judgments (Stocker and Simoncelli, 2008; Luu and Stocker, 2018), value judgments (Brehm, 1956; Festinger, 1957; Gerard and White, 1983; Steele et al., 1993; Heine and Lehman, 1997; Koster et al., 2015), and metamemory (Koriat, 2012; Zawadzka et al., 2016), suggesting it may be a domain-general strategy that serves also to reduce cognitive dissonance and improve self-consistency.

Using the absolute strength of decision-congruent evidence to judge confidence could also indicate that a confidence judgment attempts to infer the possible ***cause*** of the signals that led to the perceptual inference as externally- or internally-generated (Körding and Tenenbaum, 2007; Wei and Körding, 2011): Are these signals strong enough to indicate an external stimulus, or are they likely to simply reflect internal noise fluctuations? A mechanism that keeps track of the absolute amount of evidence, regardless of the balance, would be critical to successfully solving such a causal inference problem by allowing the system to differentiate between strong versus weak signals even when the signals themselves are equally ambiguous (i.e., equally favor one versus another possible stimulus identity). And finally, that ‘detection’ circuitry might contribute to metacognitive judgments is also supported by reports of neurons coding for the detectability (or lack thereof) of a stimulus in prefrontal cortex (Merten and Nieder, 2012), an area known to be involved in metacognitive computations (including judgments of ‘visibility, i.e. awareness) in perception and memory (Janowsky et al., 1989; Schnyer et al., 2004; Kao et al., 2005; Pannu and Kaszniak, 2005; Lau and Passingham, 2006; Fleming et al., 2010, 2012; Lau and Rosenthal, 2011; Fleming and Dolan, 2012; Middlebrooks and Sommer, 2012; McCurdy et al., 2013; Fleming and Lau, 2014).

Here we demonstrated that normalization tuning provides a biologically plausible mechanism for implementing confidence computations that demonstrate an over-reliance on decision-congruent information. Our findings lead to testable hypotheses about the role of tuned normalization in a neuron’s contribution to a decision versus a confidence judgment: activity of more normalized units should reflect an observer’s objective decisions more than confidence judgments, while the opposite should be true for less normalized neurons. Future electrophysiological studies should explore the extent to which this hypothesis can be verified in the neurobiology of perceptual decision-making circuitry. It has also been reported that tuned normalization is spatially ‘clumped’, i.e. that nearby neurons have more similar normalization profiles than neurons separated by longer distances (Ruff et al., 2016). The present findings thus pave the way for noninvasive neuroscience techniques, such as spatially coarser functional MRI in humans, to clarify the role of normalization tuning in perceptual and cognitive decisions and metacognitive evaluations of these choices.

## Author contributions

M.A.K.P. and H.L. conceived of the model; M.A.K.P. and B.M. formally specified the model and conducted model simulations; P.G., S.H.C., and M.A.B. collected the monkey data; M.A.K.P., B.O., P.G. and S.H.C. analyzed the monkey data; M.A.K.P., B.M., and H.L. wrote the manuscript.

## Acknowledgements

This work was partially supported by the National Institutes of Health (R01NS088628-01 to HL; R01EY13692 to MAB) and the Air Force Office of Scientific Research (FA-9550-15-1-0110 to HL). We thank Bosco Tjan for seminal discussion on the theoretical foundations for this project.

## Footnotes

1 We compared the simulation effect size to the average within-subject effect size in the empirical data, rather than the between-subject effect size, because the simulation effect size is computed as the difference in the mean confidence scaled by the pooled across-trial standard deviation of confidence. Thus, the simulation is effectively a simulation of a single subject who exhibits behavior resembling the across-subject average in empirical data, and the effect size computed for the simulated data is effectively a within-subject effect size that should be compared to empirical within-subject effect sizes.

